# Reconstructing substitution histories on phylogenies, with accuracy, precision, and coverage

**DOI:** 10.64898/2025.12.21.695861

**Authors:** Jordan Douglas, Lindell Bromham

**Affiliations:** Macroevolution and Macroecology, Research School of Biology, Australian National University, Canberra, Australia; Department of Physics, The University of Auckland, Auckland, New Zealand

**Keywords:** stochastic mapping, ancestral sequence reconstruction, phylogenetics, molecular evolution, synonymous substitutions, mammals, aminoacyl-tRNA synthetases, influenza, phylodynamics, beast 2, starbeast3, indels, multispecies coalescent

## Abstract

Ancestral sequences and substitution histories are usually averaged out in phylogenetic inference and are therefore not reported to the user. However, they can be recovered through ancestral sequence reconstruction (ASR) and stochastic mapping. By implementing and validating a new ASR and stochastic mapping package compatible with both single-locus and multispecies coalescent analysis, we show how reconstructing substitutions along tree branches provides a practical approach to phylogenetic inference that has a number of advantages, in terms of both efficiency and in the information gained about the tempo and mode of molecular evolution. Based on a range of simulated datasets, we observe that substitution histories are recovered more accurately and precisely on time trees with relaxed clocks compared with unconstrained substitution trees that lack temporal directionality (maximum likelihood and Bayesian). We show that codon-partition models with site-rate heterogeneity (i.e., with four nucleotide states) can effectively approximate synonymous and non-synonymous substitution histories while requiring far less runtime than computationally demanding 61-state codon models. In turn, this provides a low-bias estimator of dN/dS that can outperform existing stochastic-mapping methods. Lastly, we ground-truth the stochastic mapping approach by showing that it can recover expected patterns in molecular evolution and pathogen transmission in three different case studies: i) that smaller mammals tend to have faster sub-stitution rates than their larger relatives, ii) that the rate of change in 3Di structural-alphabet characters in the aminoacyl-tRNA synthetase anticodon binding domain is associated with amino acid substitution rate, and iii) that influenza A virus spread more frequently to nearby locations than distant ones, in a major H3N2 outbreak in New Zealand. These three effects were more pronounced when based on count estimates (via stochastic mapping) rather than evolutionary rates and branch lengths (the standard approach). Our open-source code comes with a user-friendly graphical interface, and is released as the BeastMap package for BEAST 2. The implementation is directly integrated into Bayesian phylogenetic analysis; supporting a wide range of clock, site, and tree models and data types, including insertions and deletions.

## Introduction

Comparative analysis of molecular sequences (DNA, RNA, protein) provides a universal framework for examining relationships at all levels of biological organisation, from genes to individuals, populations, and lineages. One major advantage of molecular phylogenetics is the ability to estimate a timescale for diversification. The evolutionary timescale is often the explicit focus of phylogenetic studies, for example putting dates on lineage origins, major radiations, or evolutionary innovations. But time-scaling is also implicit in a wide range of phylogenetic analyses even when absolute dates of divergence are not the primary focus, such as exploration of changes in diversification rate in time, space, and lineages (Nee et al., 1992; Liow et al., 2023; Morlon et al., 2024). The reliability of these macroevolutionary methods depends not simply on getting the relationships between taxa right -– the topology of the tree -– but critically on the accuracy of branch length construction (Duchêne et al., 2017; Etienne et al., 2016; Title and Rabosky, 2017). Branch lengths present a different kind of challenge compared to inferring relationships, and in many ways, branch lengths are a much more challenging problem to solve than topology. Different methods, or changing parameter values within the same model, can lead to quite different branch length estimates for the same topology (Brown et al., 2010; Schwartz and Mueller, 2010), impacting estimates of dates of divergence, macroevolutionary patterns of diversification, and even influencing decision-making for conservation prioritisation (Ritchie et al., 2021; Kearns et al., 2026).

Phylogenetic branch lengths have three components: dates (time duration), rates (amount of change per unit time) and distances (amount of change). If we know two of these quantities, we can estimate the third, but typically we don’t know any of them with any certainty, leading to a problem of non-identifiability. Much of the development of increasingly sophisticated methods of molecular phylogenetics has focused on ways of bracketing possible values of dates and rates to allow a range of plausible phylogenetic solutions (Bromham et al., 2018). For dates, this has led to a focus on strategies for incorporating empirical evidence of dates for some nodes in the tree to anchor the estimates of rates and dates for all other nodes, “calibrating” the molecular clock using known dates for tips, internal nodes or the root using temporal information derived from fossils, geographic distribution or sample dates (Ho and Phillips, 2009; Rieux and Balloux, 2016; Sauquet et al., 2012). In contrast to dates, the strategy for rates has been to employ statistical models that, rather than being grounded in observation, are chosen as convenient means to allow rates to vary between branches of the phylogeny. One such example is the uncorrelated relaxed clock model (Drummond et al., 2006), which assumes independence in the evolutionary rate among branches. There have been parallel efforts in allowing for different rates of accumulation of change across sites (Arenas, 2015). But despite these advances, there has been less interest in the problem of inferring the amount of change along branches than there has been in the estimation of dates and rates.

There are two broad ways to express branch lengths of a phylogeny — in terms of substitutions (amount of change) or time (absolute or relative time units). In a substitution tree, branches are expressed in units of substitutions (per site, usually), and the tree might be rooted or unrooted. Substitution trees are a common means of phylogenetic inference within maximum likelihood settings (e.g., IQ-TREE; Minh et al. (2020)) as well as some Bayesian settings (e.g., PhyloBayes; Lartillot et al. (2009)). Substitution trees can be non-ultrametric, meaning that they are not scaled to time and the tips are not constrained to line up at the present. In a time-scaled tree, or simply *time tree*, branch lengths are expressed in units of time, the tree has a direction (i.e. rooted). Time trees are commonly ultrametric, so that the tips of the tree are constrained by their sample dates which are typically in the present (but may be older for tip-dated trees). The temporal information at the tips of the phylogeny can be used to constrain the branch lengths. Time trees are the norm in Bayesian phylogenetic inference, including BEAST X (Baele et al., 2025), BEAST 2 (Bouckaert et al., 2019), and RevBayes (H öhna et al., 2016a). Expressing branch lengths as time durations relies not only on inferring the number of substitutions that have occurred, but also requires an additional suite of user-specified assumptions that enable the inference of dates and rates. In a Bayesian context these are variously referred to as models or priors, specifically a “clock model” (or branch rate prior) which describes the relationship between rates and dates, and the “tree prior” (like birth-death and coalescent models) that describes the distribution of branch lengths defined by branching or coalescent events (Bromham et al., 2018). The choice of clock model and tree prior can have major impacts on the estimated phylogeny (Wertheim et al., 2010; Ritchie et al., 2017; Douglas et al., 2025), yet we rarely have good empirical reasons for favouring one form of prior over another.

These two approaches to inferring phylogenetic branch lengths – either scaled to amount of change (in counts of substitutions) or scaled to time (as substitutions per site per unit time) – are conceptually interchangeable, given sufficient information, however in practice time trees are more readily converted to substitution trees than the reverse. But for some applications we may primarily care about amount of change. We may be interested in the way tempo and mode of molecular evolution varies over the phylogeny (Singh et al., 2009), for example through accumulation of substitutions in genes associated with environment or niche (Guevara et al., 2021). Or we might wish to contrast the relative numbers of changes in different categories, such as synonymous and non-synonymous substitutions in proteincoding genes (Ohta, 1995), or different classes of amino acid substitutions in peptides (Tourasse and Li, 2000). Or we may be primarily concerned at reconstructing the series of evolutionary changes that have occured along a phylogeny, and inferring historical states of genes, proteins or other traits (Severino et al., 2025).

Here we apply a stochastic mapping approach to estimating the number of molecular changes along edges of a phylogeny, producing branch lengths as counts of substitutions (Fig. 1). This approach is appropriate for both time trees and substitution trees. Stochastic mapping generates explicit realisations of “mutational histories” under probabilistic models, conditioned on a phylogeny and associated parameters (Nielsen, 2002; Hobolth and Stone, 2009; Irvahn and Minin, 2014). Although it has been used in several key papers, for clarity we avoid the term “mutation histories” in favour of “substitution histories”, because most mutations that occur in the history of a lineage are lost and are not reconstructed on phylogenies. Several implementations of stochastic mapping exist, in the form of *post hoc* tools like sMap (Bianchini and S ánchez-Baracaldo, 2021), and integrated within inference engines like BEAST X (Lemey et al., 2012) and RevBayes. We implement stochastic mapping as the BeastMap package for BEAST 2 (available on GitHub at http://github.com/jordandouglas/BeastMap).

**Fig. 1:**
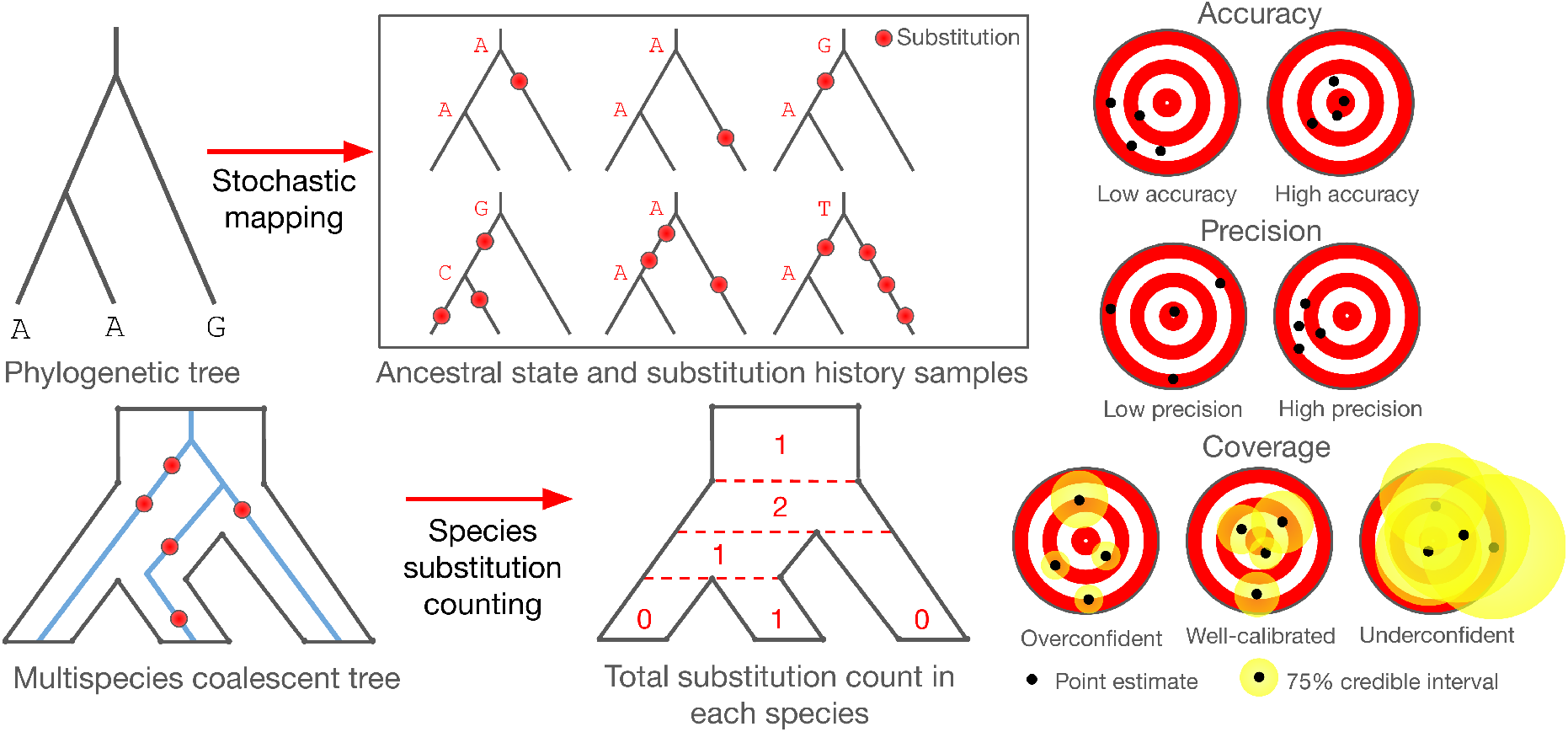
Left: using stochastic mapping, the ancestral sequences/states and substitution pathways can be sampled given a phylogenetic model. This process is more complex in the case of the multispecies coalescent, where one or more locus trees (blue) are embedded within a species tree (black). In this case, the total number of substitutions in a species is tallied up across all loci within that species (only one locus is depicted here). Right: a good phylogenetic method will have high accuracy, high precision, and achieve coverage when estimating any given quantity (i.e., substitution count in this case). These terms are readily evaluated on simulated datasets where we know the true answer. Accuracy describes how close an estimate lies, on average, to the truth (depicted by the bullseye of the dartboard). Precision describes how close these estimates lie to each other. Coverage describes whether the *p*-credible-interval (that is, a range of values that contains the true value with probability *p*) will indeed include the truth with probability *p* (in this example *p* = 0.75).

To the best of our knowledge, this is the first stochastic mapping package compatible with the multi-species coalescent. Development of the multispecies coalescent model has placed date estimation in the context of independent evolutionary histories embedded within a shared species tree (Heled and Drummond, 2009), constraining the upper-age-limit of each speciation event to be the youngest coalescence event among all loci from ancestral species. Failure to account for this process of incomplete lineage sorting can result in inaccurate estimates of divergence ages and substitution rates (Ogilvie et al., 2016, 2017; Mendes and Hahn, 2016, 2017), as well as leading to errors in topology (Kubatko and Degnan, 2007; Degnan and Rosenberg, 2009; Ogilvie et al., 2016; Heled and Drummond, 2009).

Expressing substitutions as counts is apt given the cumulative process of evolutionary change. Count data has a number of important properties that are relevant to phylogenetic inference. It is the absolute number of substitutions that provides explanatory power in evolutionary inference, not the rates or dates. Inference of branch lengths is challenging at the “shallow end”, when small numbers of observable differences between sequences lead to large errors in estimation of branch lengths, rates or dates (branch lengths are also challenging to estimate at the “deep end” where rounds of multiple hits have obscured evolutionary signal, i.e., saturation). The number of substitutions can be increased by data selection – more or longer sequences, sequences with faster rates of change, or deeper divergences that have accumulated more substitutions — but if the available sequence data is fixed then the number of substitutions places the limits on branch length estimation (Steel and Penny, 2000). In this sense, estimating the number of substitutions along branches provides a valuable appraisal of the evolutionary information in the available sequences.

This approach also provides a means of inferring ancestral sequences at every node in the phylogeny. Ancestral sequence reconstruction is generally skipped over in maximum likelihood and Bayesian approaches, which average over all possible states at the internal nodes of the phylogeny. However, sometimes we may be interested in constructing the series of steps in biochemical evolution (Eck and Dayhoff, 1966; Bridgham et al., 2009). For example, ancestral sequence/state reconstruction (ASR) of protein sequences can provide the basis for synthesis of putative past proteins and exploration of novel peptides (Gumulya and Gillam, 2017). Ancestral state reconstruction has also proven useful for the study of infectious diseases, by pinpointing the times at which pathogens have spread from one geographical location to another (Müller et al., 2018; Attwood et al., 2022; Paredes et al., 2024).

In this study we present a new, flexible implementation of stochastic mapping with a user-friendly graphical interface; all within the popular BEAST 2 platform among *>* 50 other modular packages (Bouckaert et al., 2019). This enables its interoperability with a wide range of existing options for phylogenetic inference, including a range of clock, tree, and substitution models, as well as with different discrete character data types. The implementation also allows modelling of insertions and deletions (indels) when reconstructing substitution histories. Stochastic mapping can be readily extended to different classes of change, including nucleotide, amino acid and structural characters. In particular, we show that our stochastic mapper provides an efficient and user-friendly way to estimate number of non-synonymous and synonymous substitutions.

In this paper, we first validate the stochastic mapping approach using simulated data. Then, we explore how stochastic mapping can evaluate various phylogenetic methods for their abilities to recover substitution histories from simulated datasets. Lastly, we illustrate the utility of this approach on biological systems – mammalian body sizes, coding enzyme structures, and influenza transmission. These diverse case studies provides a ground-truthing to our substitution count estimation framework, because they demonstrate that the stochastic mapper BeastMap can recover empirically observed phenomena.

## Methods

### Preliminaries

Let 𝒯 be a binary rooted tree with *n* leaf nodes (also known as taxa) and *n*− 1 internal nodes. Its branch lengths may be in (relative) time units, in which case 𝒯 is a *time tree*, or in units of substitutions per site, in which case 𝒯 is a *substitution tree*. Unless specified otherwise, our formulations will cover both scenarios. The tree 𝒯 and evolutionary parameters Θ are estimated in a Bayesian setting using Markov chain Monte Carlo (MCMC) to sample the posterior distribution: binary rooted tree with n leaf nodes (also

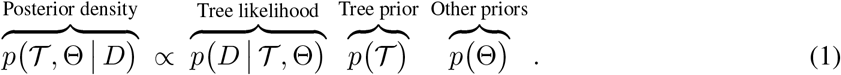

The tree likelihood follows a continuous time Markov process and is calculated using the tree-pruning algorithm (Felsenstein, 1981), while the tree prior describes a branching process, such as birth-death or coalescent in the case of time trees, or a gamma distributed branch length prior for substitution trees. The posterior distribution of trees can be summarised into a point-estimate, here using the CCD-0 method (Berling et al., 2025).

Using stochastic mapping, the substitution histories ℋ are sampled from the posterior distribution (i.e., they are not estimated as parameters):

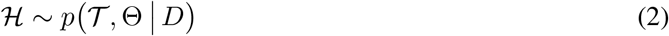

### Stochastic mapping

Substitutions are assumed to occur as independent continuous time Markov processes, where the number of substitutions over time *t* follows a Poisson(*rt*) distribution for rate *r*. These substitutions are governed by a site model, which consists of a substitution matrix and site rate heterogeneity component (where site rates are categorised into discrete bins; Soubrier et al. (2012)).

The substitutions along each branch are estimated using stochastic mapping. This algorithm has been well-studied and developed in earlier studies (Nielsen, 2002; Irvahn and Minin, 2014; Bianchini and S ánchez-Baracaldo, 2021), so here we describe it only in brief. Our stochastic mapper is implemented as a logger within BEAST 2.7.8 (Bouckaert et al., 2019), as a standalone package named BeastMap. At each sample point during MCMC (every 10,000 steps in this study), the stochastic mapper first samples the internal node sequences based on the sequences at the leaves and the current state (tree, clock, and site model); and then samples one possible substitution pathway along each branch in the tree conditional on the sequence at either end point (Fig. 1). Summary statistics of the substitution history are reported to the user, including the number of substitutions along each lineage. This implementation accounts for site rate heterogeneity by sampling the rate category of each site from their posterior probabilities prior to each round of ancestral sequence reconstruction. In the case of the multispecies coalescent (within the StarBeast3 package v1.2.1 of BEAST 2; Douglas et al. (2022)), the substitutions within each species lineage are tallied across all locus tree lineages embedded within that species. To improve computational efficiency during stochastic mapping, we employed a modified rejection sampler (Nielsen, 2002; Hobolth and Stone, 2009). Although there are more advanced methods such as direct sampling and uniformisation, they are not necessarily more efficient than the modified rejection sampler (Hobolth and Stone, 2009; Hobolth, 2008). Because substitution histories are drawn from the posterior distribution during MCMC, the stochastic mapper accounts for uncertainty in the tree, the site and substitution model, the clock model, and of course the stochastic nature of substitution histories.

### Approximating dN/dS with a nucleotide representation

Stochastic mapping can be used to estimate different classes of substitution, for example non-synonymous and synonymous substitutions in protein-coding sequences. The ratio between non-synonymous and synonymous changes is often expressed as 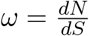, which exists as a parameter within a codon substitution model (Goldman and Yang, 1994; Yang et al., 2000). However, this approach is computationally demanding as it involves likelihood calculation under a large 61 × 61 substitution matrix. Therefore, we will estimate the number of non-synonymous and synonymous changes *nN* and *nS* under 4 × 4 nucleotide substitution matrices; conditional on the data *D*, phylogeny, and site model parameters. Paths through stop codons are allowed and are counted as non-synonymous. We describe a simple and straightforward way of estimating 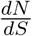 from these terms:

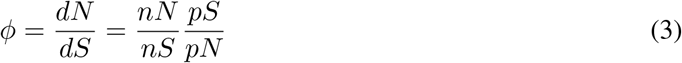

where *pN* and *pS* are the possible number of non-synonymous and synonymous substitutions, under an assumption of all substitutions being equally likely. In the standard genetic code, 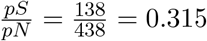 While there are much more intricate ways of approximating this term, such as an empirical Bayes method (Lemey et al., 2012), we will demonstrate that our approach provides a low-bias estimator of *ω*, and can be computed without any knowledge of nucleotide frequencies or codon site rates. Moreover, in contrast to *ω*, which is embedded within the phylogenetic model, *ϕ* is readily estimated at any level of granularity (per lineage, per clade, per site, per partition, etc.) without requiring a specialised model.

### Insertions and deletions

Indels, denoted by the gap character ‘–’ in an alignment, are typically treated as equivalent to missing information, and are omitted from the likelihood calculation. This assumption is thought to be benign during phylogenetic inference (Truszkowski and Goldman, 2016; Machado et al., 2021). When reconstructing substitution histories, however, it is important to handle indels appropriately, or the number of point-substitutions could be miscounted. In most of our analyses we are working with gap-free alignments and therefore do not need to address this issue. However our second case study (aminoacyl-tRNA synthetases) contains many gaps in the alignment. To handle these gaps, we employ an evolutionary indel model in the same style as FastML (Ashkenazy et al., 2012). This algorithm first (copies and) converts the original multiple sequence alignment *D* (data type: nucleotide, amino acids, etc.) into the simple indel coding representation *D*′ (data type: binary) described by Simmons and Ochoterena (2000). This representation does not treat each gap as an independent site, but rather it joins together contiguous gapped positions into the same indel event. Then, the gap likelihood *p*(*D*′| 𝒯) is assumed to follow a continuous time Markov process under a binary Mk substitution model (Lewis, 2001) with four gamma rate categories. ASR on the nodes of 𝒯 is performed using the compact data matrix *D*′ under this likelihood function. Lastly, the sequence reconstructions under *D*′ are back-transformed into the full size data matrix, and those sites are masked with a gap symbol when counting the number of substitutions along each lineage. Note that performing ASR with indels does not require the indel likelihood to be included in the phylogenetic model, in which case the indel site model parameters would not be estimated during MCMC, and the user would be required to provide their own estimates for the stochastic mapping step. These indel site model parameters are i) the shape parameter for gamma site rate heterogeneity and ii) its relative evolutionary rate (i.e., clock rate).

### Gamma-length and midpoint priors for substitution trees

Using stochastic mapping, we aimed to compare time trees and substitution trees for their abilities to recover substitution histories. BEAST 2 already contains a wide range of time tree models used for divergence dating, but no substitution trees. Therefore, we describe here a prior distribution on a rooted substitution tree 𝒯 based on the widely used midpoint rooting method. The midpoint of an unrooted tree is defined as the halfway point along the longest path between any two leaves. Although midpoint rooting comes with limitations (as it is sensitive to which pair of taxa are the farthest apart, and does not take into consideration sampling density), the approach is appropriate here given that these trees do not have an inherent direction of time, and has been shown to give similar results to other methods such as outgroup rooting (Hess and De Moraes Russo, 2007). In the gamma-length prior construction, branch lengths are independent and identically drawn from a gamma distribution with shape *κ* and a mean of *κβ*; in a similar fashion to PhyloBayes (Lartillot et al., 2009). The root of the tree is assumed to lie approximately halfway along the longest path between any two taxa, or more precisely 0 < *F* < 1 of the way, where *F* ∼ Beta(*α, α*) for some *α* ≥ 1. When *α* = 1, the root can lie anywhere along the longest path with equal probability; and as *α* → ∞, the root approaches the midpoint. For our purposes, we will fix *α* = 50, meaning that *F* has a 95% interquartile range of (0.4, 0.6). However, as we are using time-reversible substitution models and are not incorporating any divergence date calibrations, there will be no information that informs root placement in this study, beyond the decision to fix *α* = 50. The gamma-length midpoint tree model is laid out more formally in Supporting Information Section 1.

### Summary statistics for simulated data

We can benchmark methods for their ability to recover the trees 𝒯, parameters Θ, and substitution histories ℋ, using simulated data where the true values are known. Let **x** be a vector of known quantities where *x*_*i*_ ≥ 0 (e.g., root-to-tip distances; evolutionary rates; or substitution counts per lineage or per tree), and let ***ŷ*** be estimates of the term sampled from a posterior distribution, where (***ŷ***_**0**_, ***ŷ***_**1**_, ***ŷ***_**2**_) respectively denote the lower, median, and upper credible intervals of **y**. In this case, the credible interval [***ŷ***_**0**_, ***ŷ***_**2**_] is the 95% highest posterior density (HPD) interval. **x** is generated by simulating multiple sequence alignments *D*, making sure to keep track substitution histories ℋ along the way (via the Gillespie algorithm; Gillespie (1977)). **y** is generated by performing inference on *D* under a phylogenetic model (via MCMC). A statistical estimator, in this case a phylogenetic inference method, can then be evaluated using (Fig. 1):

1. **Accuracy:** measured as the relative error of the point estimate. When *x*_*i*_ = 0, we will add a pseudocount of 0.1 (substitutions) to avoid dividing by zero.

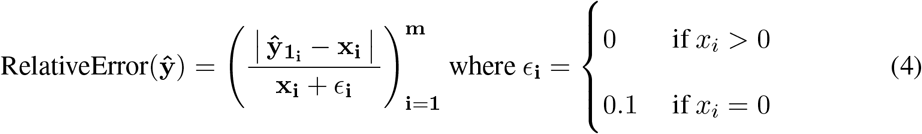

To diminish the potential impact of extreme values and outliers, we sorted all estimates **ŷ** in increasing order of relative error and discarded the top 5%, for all estimators. The mean relative error (a measure of in-accuracy) is calculated as:

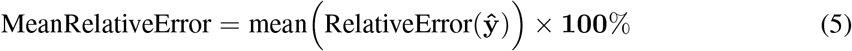
2. **Precision:** measured as the standard deviation of the relative error (with smaller values having greater precision). Just like accuracy, the top 5% of extreme error values were filtered out in each estimator.

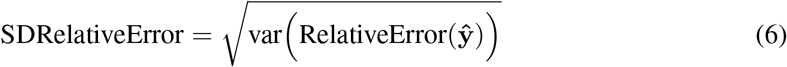
3. **Coverage:** the proportion of true values that are contained within the 95% HPD. If an estimator is valid, then its coverage will be 95%. Having coverage close to 95% (which comes to 91-99% within the sampling error of *m* = 100 replicates) means that we can rely on the estimator’s level of confidence. However, if the coverage is too high (e.g., 100%), then the method is uninformative (achieving 100% coverage can be obtained by any estimator by setting the credible interval to (−∞, ∞)).

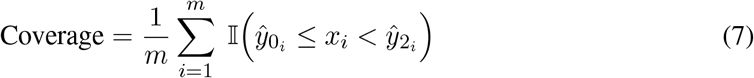
4. **Correlation:** the Pearson correlation *ρ* measures the agreement between the true values **x** and their point estimates **ŷ**_**1**_.

A good estimator should score well on all four metrics. Accuracy describes how close the point estimate is from the truth (low error mean), while precision describes how close estimates are to each other (low error variance). These two metrics are both based on point estimates, rather than the full posterior distribution, in order to enable a fair comparison across estimators that may give vastly different posterior distribution variances. Within a Bayesian perspective, having coverage close to 95% (which comes to 91-99% within the sampling error of *m* = 100 observations) means that we can rely on the estimator’s level of confidence. A high positive correlation between the true and estimated value indicates that we can learn something from the data and we are not just sampling from the prior distribution.

### Substitution count estimators

Our stochastic mapper is implemented in BEAST 2 with a user-friendly interface so can be used in combination with a wide range of methods available in BEAST 2. To illustrate the application of the stochastic mapper, and to compare its behaviour in different phylogenetic inference methods, we compare four methods for their abilities to estimate substitution counts from a multiple sequence alignment that was simulated under known conditions. Three of these estimators are Bayesian phylogenetic methods implemented in BEAST 2 – the gamma-length / midpoint tree prior described here (GM), the birth-death tree prior with a strict clock (BD-S; constant clock rate on all edges), and the birth-death tree prior with a relaxed clock (BD-R; under the ORC model v1.2.1 (Douglas et al., 2021)). The stochastic mapper is applied during MCMC, which captures uncertainty behind the tree, site model, and clock model; as well as the substitution histories themselves. The fourth method is a maximum likelihood (ML) tree estimated by IQ-TREE v1.6.12 (Minh et al., 2020) and rooted using the midpoint method. To estimate substitution histories on an ML tree, we ran our stochastic sampler 200 times on the ML estimates of the respective tree and site model parameters. In all cases, inference was performed under the `HKY+G4+F` site model, with the `-gmedian` option enabled in IQ-TREE so that the gamma rate heterogeneity implementation matches the one used in BEAST 2. These four methods are explicated further in the Supporting Information.

### Benchmark datasets

To assess substitution history estimators, we simulated 400 datasets, evenly distributed across four sets of conditions, which are used to benchmark the estimators. In each case, *n* = 40 nucleotide sequences with 300 sites were simulated under an HKY substitution model with non-uniform frequencies, and four gamma rate categories. The selected speciation and extinction rate parameters resulted in trees with around 0.5 substitutions per site from root to tip, describing the case where sequences have undergone significant change, but not to the point of saturation. Simulations were performed using BEAST 2 and the TESS package for R (H öhna et al., 2016b). These datasets are summarised on the top of Fig. 2 and briefly described below, with further information found in Supporting Information.

**Fig. 2:**
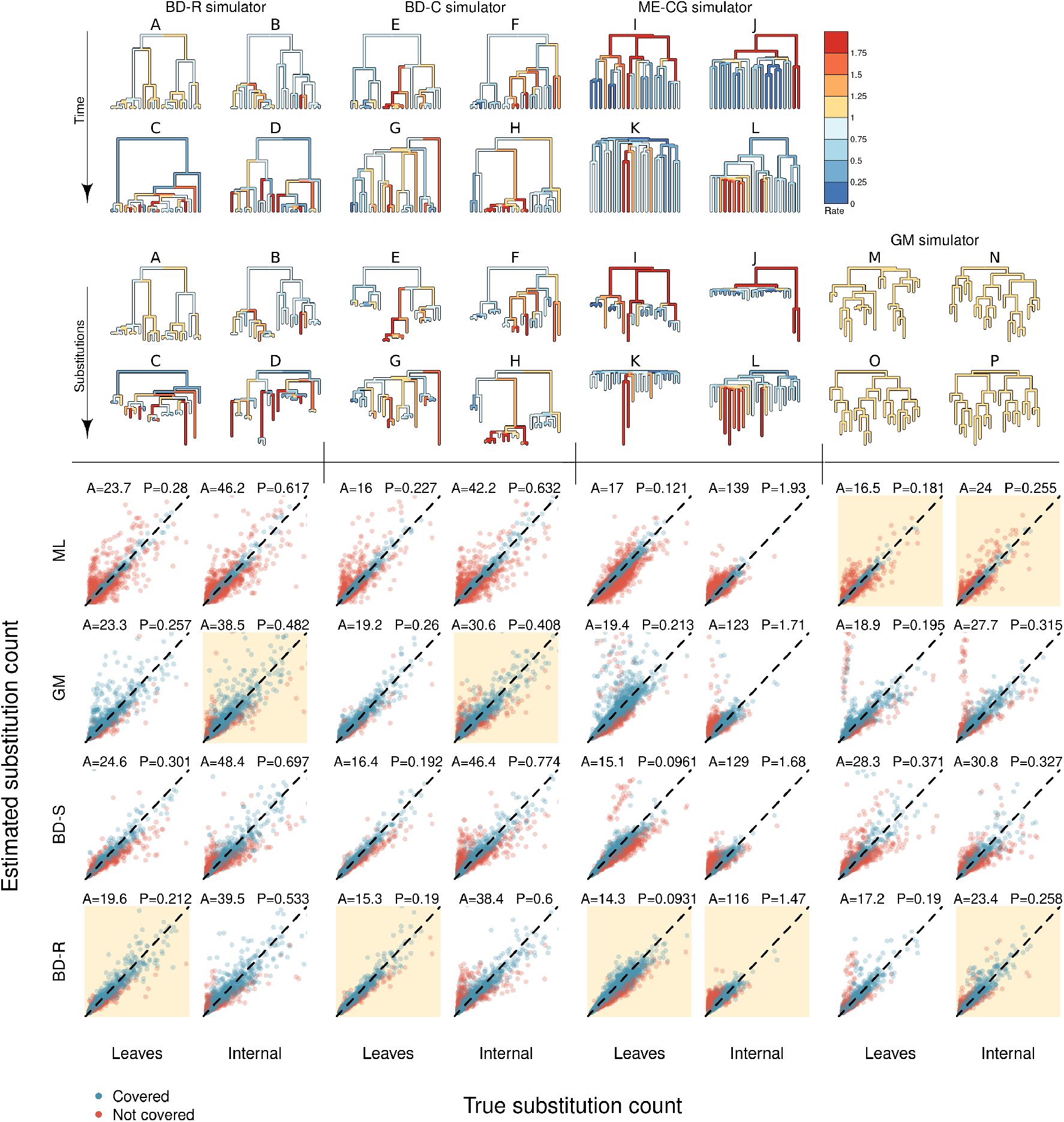
Top: four methods were used for simulating datasets, each depicted here on compact 20-taxon trees. These time trees are coloured by relative branch rates, and their equivalent substitution trees describe the expected number of substitutions along each lineage. The exemplar BD-R trees A–D have increasing levels of variability *σ* = (0.1, 0.3, 0.6, 0.8). The exemplar BD-C trees I and J have a clock rate that decreases through time, while K and L are increasing. Bottom: based 100 datasets from each simulator, the accuracy and precision is denoted above each plot, with high accuracy measured by small A (mean relative error, %) and high precision measured by small P (standard deviation of relative error). The estimator that achieved the best accuracy or precision (usually both) for a given simulator is highlighted in gold. Points are coloured blue when the true value lies within the 95% credible interval, and orange if not. Overall, the BD-R estimator outperformed in most cases, while ML trees were overconfident (i.e., low coverage). We refer the reader to Tables S1–S4 for more detailed results.

#### BD-R

The **BD-R** datasets were simulated using a birth-death (**BD**) tree (Gernhard, 2008) and an uncorrelated relaxed (**R**) clock (Drummond et al., 2006; Douglas et al., 2021). These time trees have sequences that range from very clock-like to very non-clock-like (quantified by the branch rate standard deviation). In this clock model, lineages are assigned relative substitution rates from a log-normal distribution, with a mean-in-real-space of 1 and a standard deviation *σ*, where *σ* ∼ Uniform(0, 1). As *σ* approaches 0, the rate variation becomes smaller and the sequences become more clock-like. When *σ* ≳ 0.8, the sequences strongly violate the assumption of constant evolutionary rate of change enforced by strict clocks. A typical multispecies dataset has *σ* ranging from 0.2 to 0.5 Douglas et al. (2021).

#### BD-C

The **BD-C** datasets were also simulated under birth-death time trees, however the substitution rates were correlated across branches (Drummond and Suchard, 2010). In this clock model, each lineage inherits a clock rate from its parent, with some variability. Given that many of the influences on average rate of molecular evolution are heritable (e.g., body size), we expect substitution rates to be inherited from the parent lineage with modification, so the correlated clock model provides a more biologically plausible scenario than the uncorrelated relaxed clock, while also violating the assumption of independent branch rates under a relaxed clock.

#### ME-CG

In order to test the performance under non-ideal conditions, the **ME-CG** datasets make two pathological departures from standard phylogenetic assumptions. First, these time trees were simulated under the assumption of two mass extinction (**ME**) events and three epochs (i.e., global shifts across all lineages) of birth-death rates *λ* and *µ*. These departures from a constant birth-death process result in longer tip edges and shorter internal edges (i.e., trees I, J, K, and L of Fig. 2). Second, the sequences were simulated with a directional bias in substitution rates (clock gradient, **CG**) where the clock rate increases or decreases through time on average, in a noisy fashion. These properties – long terminal branches and biased change in substitution rates – both challenge the assumptions made by the tree priors and clock models in the methods being compared here.

#### GM

The **GM** trees were generated under the gamma-length / midpoint prior; and sequences under a strict clock. In contrast to time tree models, the tips of the phylogeny are not fixed in time, and all branches have the same expected length. This departs from the expectation under a time-scaled tree, where the tips are constrained in time and branch lengths and node heights are distributed along a time axis. Therefore this represents a violation of the model of Bayesian phylogenetic methods that incorporate time-scale into the estimation of branch lengths.

## Results

### Validating the stochastic mapper and gamma-length tree prior

We validated our methods using coverage simulation studies (Mendes et al., 2025). This is achieved by i) sampling parameters Θ and a tree 𝒯 from a probability distribution; ii) simulating a multiple sequence alignment *D* under *θ* and 𝒯 while keeping track of the substitution histories ℋ along each branch; iii) performing MCMC on data *D* to obtain estimates 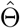, 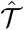, and 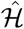; and lastly iv) comparing Θ with 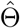, the branch times in 𝒯 with those in 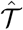, and the substitution counts in ℋ with those in 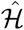. These four steps were performed one hundred times. When the model is correctly implemented, one would expect the true value of a term to reside in its 95% credible interval in approximately 95% of all replicates (in which case we say it has 95% coverage). As shown in Fig S3, these trials confirmed the usefulness and correctness of the newly implemented stochastic mapping algorithm and gamma-length prior, which yielded estimates that were well-correlated with the true values and had around 95% coverage. The prior distributions used in these simulation studies are shown in Supporting Information Section 3.1.

### Comparison of phylogenetic inference methods using stochastic mapping

We used stochastic mapping to benchmark four phylogenetic methods for their abilities to estimate substitution histories: i) a maximum likelihood substitution tree (ML), ii) a Bayesian gamma-length substitution tree rooted using the midpoint prior (GM), iii) a Bayesian strict clock birth–death tree (BD-S), and iv) a Bayesian relaxed clock birth–death tree (BD-R). Each method was tested on 400 datasets generated by four simulators, denoted by bold font, designed to stress-test performance (Fig. 2): i) a birth–death time tree with highly variable branch rates (**BD-R**), ii) a birth-death tree with a correlated clock model (**BD-C**), iii) a birth–death tree with mass extinctions and a time-varying average clock rate (**ME-CG**), and iv) a substitution tree agnostic to time (**GM**). In all cases, the inference model and priors differed from the generating model (i.e., model misspecification). Of these four settings, **ME-CG** proved the most challenging. The parameters used in these trials are shown in Supporting Information Sections 3.2 and 3.3.

In terms of coverage, these results confirmed that the Bayesian methods (GM, BD-S, and BD-R) were significantly more reliable at confidence assessment compared with the method sampling substitution histories from the ML tree, which was consistently overconfident in its estimates (Fig. 2 and Table S1–S4). Although the Bayesian methods did not always achieve 95% coverage, their performance was strongest on the **BD-R** and **GM** datasets, and less reliable on **ME-CG**. The relaxed clock BD-R achieved coverage more frequently (6/12 times) than either GM (4/12) or BD-S (1/12); while ML did not achieve coverage in any setting.

In terms of accuracy and precision, BD-R ranked first overall (Fig. 2 and Table S1–S4). BD-R achieved the lowest overall mean (i.e., high accuracy) and standard deviation (i.e., high precision) of error on the **BD-R, BD-C**, and **ME-CG** datasets. By contrast, ML performed best on the unconstrained (**GM**) datasets, and surprisingly it even outperformed the Bayesian method that used a GM prior. However, BDR was a close second in this setting, despite its assumptions of clock-like evolution and ultrametric trees being violated by these datasets. The strict clock BD-S performed the worst on **BD-R** and **GM**, while the gamma-length prior was the least reliable on **ME-CG**. The poor performance of BD-S is unsurprising, as these datasets are highly un-clock-like, but the strict clock assumes a constant rate of evolutionary change through time. In statistical inference, there is often thought to be a trade-off between accuracy and precision. In these trials however, the most accurate estimator in any given band is often the most precise.

These trials also demonstrated that there is not a clear, mutual relationship between the branch rate standard deviation *σ* (within a birth-death tree) and the branch shape parameter *κ* under the gamma-length tree prior (Fig. S1). These two parameters are negatively correlated – a high variation in branch rate under a time tree (high *σ*) is associated with a high variation in branch length under a substitution tree (low *κ*). However, the relationship between *σ* and *κ* is poorly defined. When datasets were simulated under the the **BD-R** model, as *σ* approaches 0, *κ* approaches ≈ 1. However, in the reverse direction, where the true *κ* <≈ 1, the estimated *σ* is around 0.8; corresponding to extremely non-clock-like evolution with high rate variability (Fig. 2 tree D). The gamma-length also underperformed when estimating changes on leaves and across whole phylogenies, but performed surprisingly well on internal branch substitution counts.

Overall, in these trials the uncorrelated relaxed clock model (BD-R) performed best: in the parameter spaces we explored, the BD-R trees had the highest ranking accuracy and precision in more bands than any other estimator, and achieved coverage more often too. This approach not only outperformed maximum likelihood (substitution) trees, but it also provided a more self-aware assessment of its uncertainty. The gamma-length / midpoint tree prior is also quite challenging to interpret as an evolutionary model as it does not describe a temporal process. Going forward in this study, we will consider relaxed clocks and time trees.

### Counting nN and nS gives a fast and accessible estimate of dN/dS

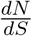 is commonly estimated as the *ω* parameter of a codon substitution matrix (Biou et al., 1994), and sometimes through stochastic mapping on codon sequences (Guéguen and Duret, 2018). However, both approaches are computationally demanding, particularly in a Bayesian context, due to the size of the codon substitution matrix (61×61). Here, we explore how performing stochastic mapping on nucleotides (with a 4 × 4 matrix) can offer a tractable alternative for 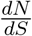 estimation.

To do this, we first benchmarked eight nucleotide models for their abilities to recover phylogenetic branch lengths and substitution counts from codon-simulated data. This was done by performing Bayesian phylogenetic inference on 100 datasets, each simulated under a codon substitution matrix with random *ω* and gamma site-rate heterogeneity. Each method was benchmarked using its coverage. Models that achieved less than 90% coverage are interpreted as overconfident and therefore unreliable in a Bayesian context, while any on 100% would be considered uninformative. The eight models varied in their use of partition model and site rate heterogeneity. The unpartitioned models (P1) applied a single site and substitution model across all sites; estimated by bModelTest (Bouckaert and Drummond, 2017), while the codon partition model (P3) assigned each codon position (1, 2, and 3) an independent site and substitution model (bModelTest), plus relative clock rate. The four site rate heterogeneity variants being compared were: i) no heterogeneity (all sites in the partition have the same rate), ii) gamma rate heterogeneity G4 (site rates are integrated over a gamma distribution binned into four categories; Yang (1996)), iii) free rates model R4 (site rates are integrated over four equally sized bins with independent rates; Soubrier et al. (2012)), and iv) free rates with free weights R4+W4 (same as R4 but the bin weights are also estimated). The parameters used in these trials are shown in Supporting Information Sections 3.4 and 3.5.

As shown in Table 1, most methods achieved much less than 90% coverage, which generally resulted from underestimating the substitution counts and branch lengths. Achieving coverage on the expected number of substitutions per site (tree height and length) proved to be a low bar – this test was much easier for the estimators to pass than substitution counts. Among the substitution counts, the total count *nS* + *nN* had the worst overall coverage, followed by *nS*, and then *nN* . This may be a reflection of how sampling error, and therefore credible interval, is more pronounced for smaller counts (*nN* ) than larger ones (*nS* and then total); as the standard deviation of a Poisson distribution increases with the square root of its expected value.

**Table 1:** Benchmarking nucleotide models for their abilities to recover known parameters from a simulated codon alignment. The coverage (%) is reported for each term above. Methods that achieved between 91 and 99% coverage (out of 100 replicates) are highlighted in bold. The tree height and length are respectively the expected number of nucleotide substitutions per site, from root to tips (height) and summed across all branches (length). Notation: P1: one partition; P3: three codon partitions (each with their own relative rate, site model, and substitution model); G4: gamma site heterogeneity with four rate categories; R4: free rates model with four estimated rate categories; W4: free rates model with four estimated category weights.

| Model | Total subst. | Total $nN$ | Total $nS$ | Tree height | Tree length |
| --- | --- | --- | --- | --- | --- |
| P1 | 8 | 34 | 18 | 36 | 33 |
| P1 + G4 | 67 | 60 | 45 | 88 | 90 |
| P1 + R4 | 71 | 66 | 46 | **92** | 88 |
| P1 + R4 + W4 | 70 | 64 | 45 | 90 | 89 |
| P3 | 13 | 32 | 19 | 46 | 38 |
| P3 + G4 | 87 | 86 | 90 | **92** | 87 |
| P3 + R4 | **92** | 90 | **92** | **93** | 90 |
| P3 + R4 + W4 | **96** | **95** | **98** | **92** | **91** |

The codon partition models generally outperformed the no-partition models, consistent with earlier findings (Shapiro et al., 2006). The free rate models (R4) generally outperformed the gamma rate (G4) models, consistent with reports that gamma rate heterogeneity makes for a rather crude description of site rate heterogeneity (Ferretti et al., 2024). Remarkably, the P3 model with no site rate heterogeneity was significantly outperformed by the unpartitioned models that did have rate heterogeneity (e.g., P1 + G4). It is noted that this site-homogeneous P3 model is employed by the Counting Renaissance method in BEAST X (Lemey et al., 2012).

To test whether nucleotide and codon models give similar outcomes in biological datasets, we estimated the per-lineage *nN* and *nS* in four mammalian loci using nucleotides (P3+R4+W4) and codons (M0+F61+R4+W4). These genes were obtained from the annotated OrthoMaM v12 database of orthologous mammalian coding regions (Allio et al., 2024) – CLDN16 (entry 10686), SERTAD3 (entry 29946), RNASE (entry 390443), and TGOLN2 (entry 10618). After removing all gapped sites and sequences with large numbers of gaps (which ensured compatibility with the BEAST 2 CodonSubstModel package (Xie and Bouckaert, 2018)) the trimmed alignments consisted of 432-618 nucleotides over 72-177 taxa. In each case, inference was done under a birth-death tree prior and a gamma spike relaxed clock model to test for punctuated evolution (Douglas et al., 2025). When this hypothesis is rejected, the clock model reverts to the uncorrelated relaxed clock.

From a runtime standpoint, the nucleotide representation was significantly favourable on these biological datasets. Even with access to a state-of-the-art CPU and GPU, the computational performance under nucleotides was still 15 – 40 times faster than codons. Nucleotide representations sampled one million states from the posterior every three to eight minutes, while the codon representations required between 85 (RNASE) and 300 (CLDN16) minutes. The fastest performance here was found using a CPU for nucleotides (AMD Ryzen Threadripper PRO 5975WX), and multithreading on a GPU for codons (NVIDIA RTX A4500), both with likelihood calculations by BEAGLE v4.0.1 (Ayres et al., 2019).

As confirmed in Fig. 3, the nucleotide representation gave quite similar estimates for *nN* and *nS* as the codons, with a mean relative difference of around 5%; for both simulated and biological data. All four mammalian loci were under purifying selection, with 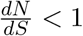. Together with the simulation studies, these results suggest that a codon-partitioned nucleotide representation appears to be a more efficient use of computational resources than codons, even if the former cannot directly estimate *ω*. The gamma spike clock model was rejected on all loci (each with less than 5% posterior support), suggesting that there was no major effect of punctuated equilibrium in these particular loci.

**Fig. 3:**
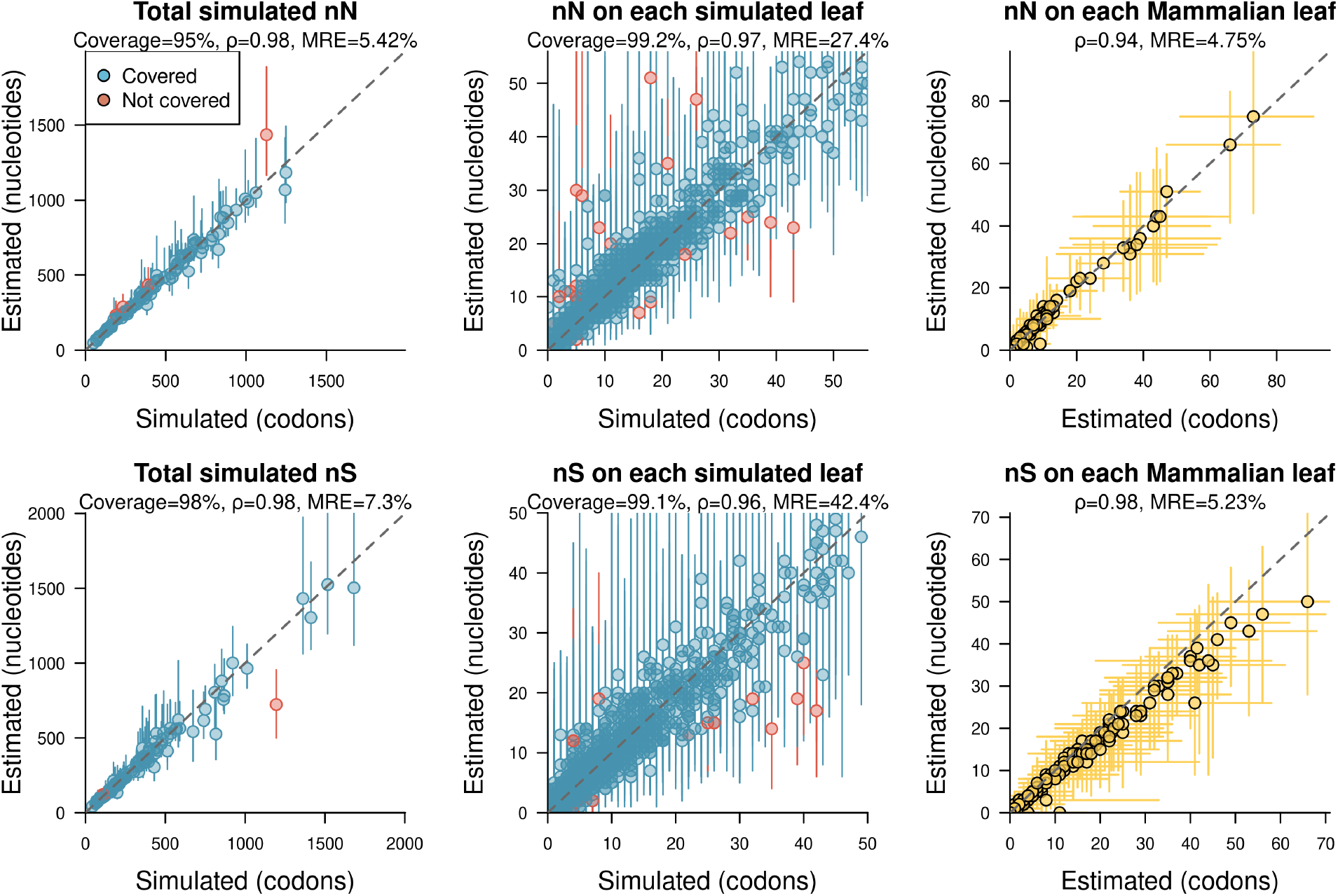
Comparison of codon (x-axis) and nucleotide (y-axis) representations in phylogenetics. We considered 100 simulated codon datasets, where the truth was known, and four Mammalian loci, where the truth was not known. The simulated alignments were sequences of 200 codons (with 61 × 61 substitution matrices and varying *ω*), and represented as alignments of 600 nucleotides during inference (with 4 × 4 substitution matrices). These trials confirmed that nucleotide models (P3+R4+W4) can accurately recover the substitution histories of codon sequences without the need for computationally demanding 61 × 61 substitution matrices. The mean relative errors (MRE) between the codon and nucleotide point values are reported; noting that “error” is a slightly misleading term in the case of the Mammalian data as we do not know the true values of *nN* or *nS*. Points represent mean estimates or true values, while vertical and horizontal bars are 95% credible intervals, for a single dataset or a single leaf.

Lastly, we benchmarked our nucleotide-based estimator for 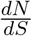 against those described by Lemey et al. (2012). Our own estimator *ϕ* is very straightforward – it is the ratio of 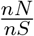 (estimated through stochastic mapping) multiplied by a constant term 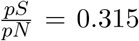. The estimators described in the 2012 study are also derived from non-synonymous and synonymous substitution counts from nucleotide analysis. Their first estimator is defined as 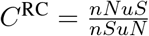, where *uN* and *uS* are sampled unconditionally on the tip data. However, as noted by Lemey et al. (2012), this method, just like our own *ϕ*, can evaluate to zero or infinity, and may be prone to high levels of stochastic noise, particularly on a per-site basis. Therefore, they introduced an empirical Bayes estimator *ω*^RC^ that was designed to estimate one 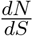 term per site. In our simulated datasets, all sites have the same value of *ω*. We tested the ability of *ω*^RC^ to estimate 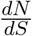 across all sites in the alignment by taking its median value across all positions. It is noted that *C*^RC^ and *ω*^RC^ are both implemented in BEAST X, but not BEAST 2, but the former implementation of stochastic mapping is not readily compatible with site rate heterogeneity. To enable a direct comparison between these methods we therefore computed all terms from the numbers logged by BEAST X.

On our simulated datasets, *ϕ* obtained the smallest relative error on average (i.e., the highest accuracy), followed by *C*^RC^, and then *ω*^RC^ (Fig. S2). Specifically, the *ϕ* estimator achieved a mean relative error of 0.271 on the simulated datasets under the P3+R4+W4 model (BEAST 2 implementation) and 0.291 under the P3 model without site heterogeneity (BEAST X implementation). These errors were both smaller than the *C*^RC^ estimator on 0.353 and *ω*^RC^ on 0.439 (both implemented in BEAST X). Based on these 100 simulated datasets, these results provide further evidence that P3+R4+W4 is more reliable than P3 without site heterogeneity, and also suggest that this rudimentary measure *ϕ* might be more accurate for full-sequence 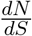 estimation than methods previously developed for single-site 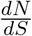 estimation.

### Case study 1: Substitution rate and body size in mammals

With simulation studies out of the way, we proceed to our empirical case studies. Each case study will serve as a means to ground-truth the stochastic mapping approach, by comparing our estimates with expected patterns of molecular evolution.

We will begin with mammals, which provide a data-rich, well-studied system with well-known patterns of molecular evolution. Although there is debate around timing of mammalian divergence events (Phillips and Fruciano, 2018; Foley et al., 2023; Liu et al., 2023), there is one aspect of mammalian molecular evolution confirmed by many independent studies: smaller-bodied species tend to have faster rates of molecular evolution than their larger-bodied relatives (Bromham, 2011). This observation was made very early in the study of molecular evolution (e.g., Ohta (1972); Li and Tanimura (1987)) and has been demonstrated for many different datasets using a range of different methods (e.g., Bromham et al. (1996); Cagan et al. (2022); Bergeron et al. (2023); Galtier et al. (2009); Zhu et al. (2025)). Therefore, although we do not know the absolute number of substitutions to expect on each branch for each locus, we do have a strong expectation that there should be comparatively more substitutions on a branch leading to a smaller-bodied species than its larger-bodied relative. Furthermore, we expect the body size effect should be more strongly evident on synonymous substitutions than non-synonymous substitutions, both on empirical grounds (Welch et al., 2008) and as a prediction of theory: if substitution rate differences are driven by underlying differences in the mutation rate, then this will be reflected in synonymous substitution rate which is largely determined by mutation rate; Bromham (2011). We can use this expected pattern to ground-truth our inference of substitution histories.

Comparing relative or absolute substitution rates between species is challenging for a number of reasons (Lanfear et al., 2010b). We cannot treat species as independent observations of the relationship between substitution rates and species traits, because related species are likely to be similar in both traits and rates. We can overcome this problem of phylogenetic non-independence of rates and traits by comparing pairs of taxa that differ in the trait of interest (in this case, body size). Since they share a common ancestor, both species in the pair start from the same initial state, and they have the same amount of time to accumulate both substitutions and changes in the trait value. As long as each pair is chosen so the path representing their shared history does not intersect any other pair’s path on a phylogeny, then each pair resembles one independent trial of an evolutionary experiment of the effect of the trait on substitution rate. With enough phylogenetically independent pairs, we can ask if change in the predictor variable tends to result in a change in the dependent variable, following the procedure characterised by Welch and Waxman (2008), and used by many others (Garland Jr et al., 1992; Bromham et al., 1996; Lanfear et al., 2010a; Ivan et al., 2022). By repeatedly sampling trees from the posterior distribution, and independent taxon pairs from each tree, we can assess the robustness of this relationship. By extending this test to multiple genes within a multispecies coalescent model, we can allow for the differences in the coalescent times for different loci to provide more accurate estimates of species branch lengths (Heled and Drummond, 2009; Ogilvie et al., 2016; Mendes and Hahn, 2016).

A number of hypotheses have been put forward to explain why smaller mammal species tend to have faster rates of substitution, but most of them rest on increase in the underlying mutation rate (e.g. due to more replication errors or lower investment in repair; Bromham (2009)). Similarly, a number of drivers of substitution rates have been suggested, including generation time, longevity and metabolic rate, but all of these life history traits scale with body size in mammals (Welch et al., 2008) so body size is an effective proxy for a range of hypotheses about the causes of substitution rate differences in mammals. To test the relationship between synonymous substitutions *nS* and body size, we estimated a phylogeny of mammal species under the multispecies coalescent based on 50 gap-free gene coding regions sourced from OrthoMaM v12 (Allio et al., 2024), which includes one sequence per sampled species, plus average adult body mass data sourced from PanTHERIA (Jones et al., 2009). We use an algorithm to select pairs of mammal species that differ in body size, within a given range of node heights (see Supplementary Information for details). To test the robustness of the conclusions to pair selection, we repeat the analysis on 10,000 sets of taxon pairs. Each replicate compared an average of *n* = 28 taxon pairs (lower: *n* = 21, upper: *n* = 36). Then, using BeastMap, we estimated *nS* on each species lineage of each pair (summed across all loci) and compared these terms with the difference in species’ adult body mass for each pair.

We observed a negative association between adult body mass and mutation rate (Fig. 4) which was significant in 68% of all replicates, at a threshold of *p* < 0.01, and significant on 31% at *p* < 0.001. By contrast, the trend was less apparent when considering total substitution count (*nN* +*nS*); coming to 59% (*p* < 0.01) and 21% (*p* < 0.001). The trend was even weaker when looking at branch distances (time duration multiplied by relaxed clock rate) rather than substitution counts: coming to 52% (*p* < 0.01) and 13% (*p* < 0.001). These results provide confidence in the stochastic mapping approach to reconstructing substitution counts.

**Fig. 4:**
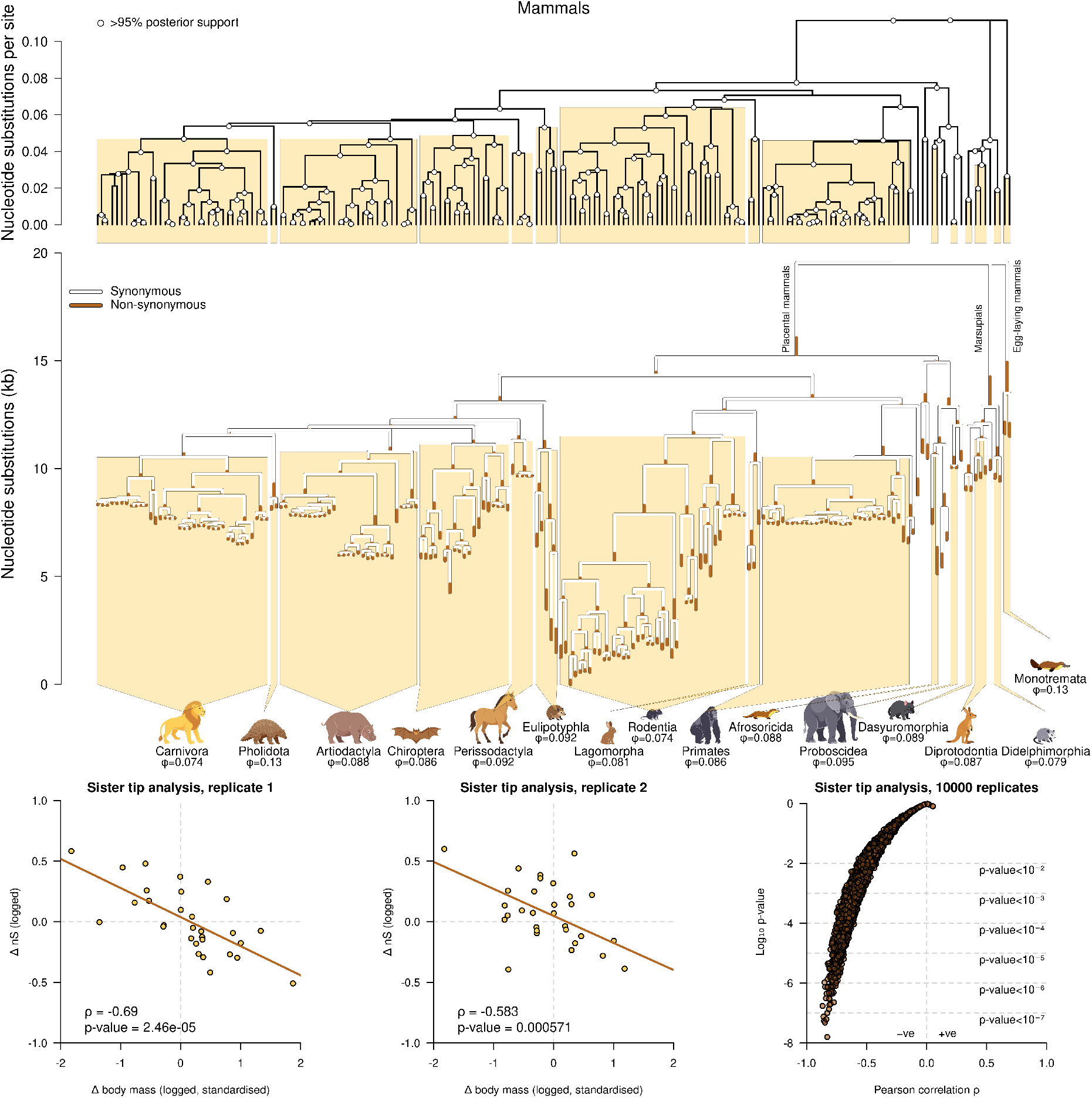
Top: mammalian species tree inferred under the multispecies coalescent with a relaxed clock model from 50 coding regions. Middle: using BeastMap, we estimated the number of synonymous and non-synonymous substitutions along each lineage (summed across all 50 loci). Shown here is the median estimate of either term on a CCD0 summary tree (same tree as above). Orders with at least two taxa are indicated along with 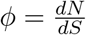 across the entire clade. Bottom: using a phylogenetic independent contrast method, we measured the relationship between the difference in synonymous substitution count *nS* and the difference in mammal body mass across independent pairs of taxa. By repeating this experiment across different random samples of taxon pairs (each with an ancestral node height from 0.01 to 0.03 substitutions per site), we observed a general negative trend between body mass and synonymous substitution rate, and usually at a statistically significant level. This is consistent with the widely observed trend that smaller mammals tend to have faster rates of molecular evolution.

### Case study 2: Association between sequence and structural change in a protein

In this case study, we demonstrate the use of stochastic mapping to estimate amounts of change in different classes of changes derived from gene sequences, amino acid sequence (aa) and three dimensional protein structures (3Di). The amino acid sequence of a protein is the main determinant of its three dimensional structure. Recent advances have enabled phylogenetic analysis of protein structures from sequences of structurally-aware characters (Puente-Lelievre et al., 2023; Moi et al., 2025; Garg and Hochberg, 2025). In this approach, protein structures are represented as linear sequences of 3Di characters (Van Kempen et al., 2024), enabling phylogenetic reconstruction in the same manner as amino acid sequences. We will test whether there is a correlation between sequence and structural change with stochastic mapping, using a tRNA anticodon binding domain as a case study.

The translation of codons in protein-coding gene sequences to amino acids is determined by the aminoacyl-tRNA synthetases (aaRS), a large group of enzymes that attach amino acids to their cognate tRNAs (Gomez and Ibba, 2020). Their catalytic domains have been classified into 37 evolutionary families based on their biological function, protein structure, and phylogeny (Douglas et al., 2024b). Eight of the these families are appended to a common anticodon binding domain (ABD) fold within the enzyme, facilitating tRNA recognition. This ABD is named CRIMVLG after its aaRS families (Douglas et al., 2024b): CysRS, ArgRS, IleRS, MetRS, ValRS, LeuRS-A and LeuRS-B (i.e., the archaeal and bacterial forms of LeuRS), and GlyRS-B (i.e., bacterial-like GlyRS). The domain exists as a helical bundle approximately 100–150 amino acids in length (Hauenstein et al., 2004; Tukalo et al., 2005).

We estimated the phylogeny of 143 CRIMVLG protein structural models from AARS Online (Douglas et al., 2024b) using three classes of characters split into independent partitions: aa sequences, 3Di characters, and indels. These structural models were predominantly AlphaFold2 predictions (Jumper et al., 2021) sourced in a taxonomically representative fashion across the tree of life, while 14 were determined by X-ray crystallography. Sequence and structural alignments were generated by FoldMason (Gilchrist et al., 2024). We used BeastMap to estimate the number of aa substitutions and 3Di structural changes on each lineage. These sequences are highly variable in length, reflecting a long history of insertions and deletions. Therefore, the use of a phylogenetic indel model allowed our ancestral reconstructions to account for this process. All three classes of characters were estimated under the same phylogenetic time tree, but with their own independent relaxed clock model; enabling their branch lengths to vary under a shared topology. One limitation here comes from the assumption that these three classes evolved independently, conditional on the phylogeny.

This analysis suggests a strong, linear relationship between the rates of aa and 3Di change, which is expected given the relationship between protein sequence and structure (Fig. 5). However, this relationship was only detected through the use of stochastic mapping (i.e., by counting the number of changes per lineage). In contrast, there was no correlation between these two rates when looking at branch rates estimated under the relaxed clock model. This suggests that, for this particular dataset at least, stochastic mapping provided a more reliable means of estimating branch rates than the actual branch-rate parameter in the relaxed clock.

**Fig. 5:**
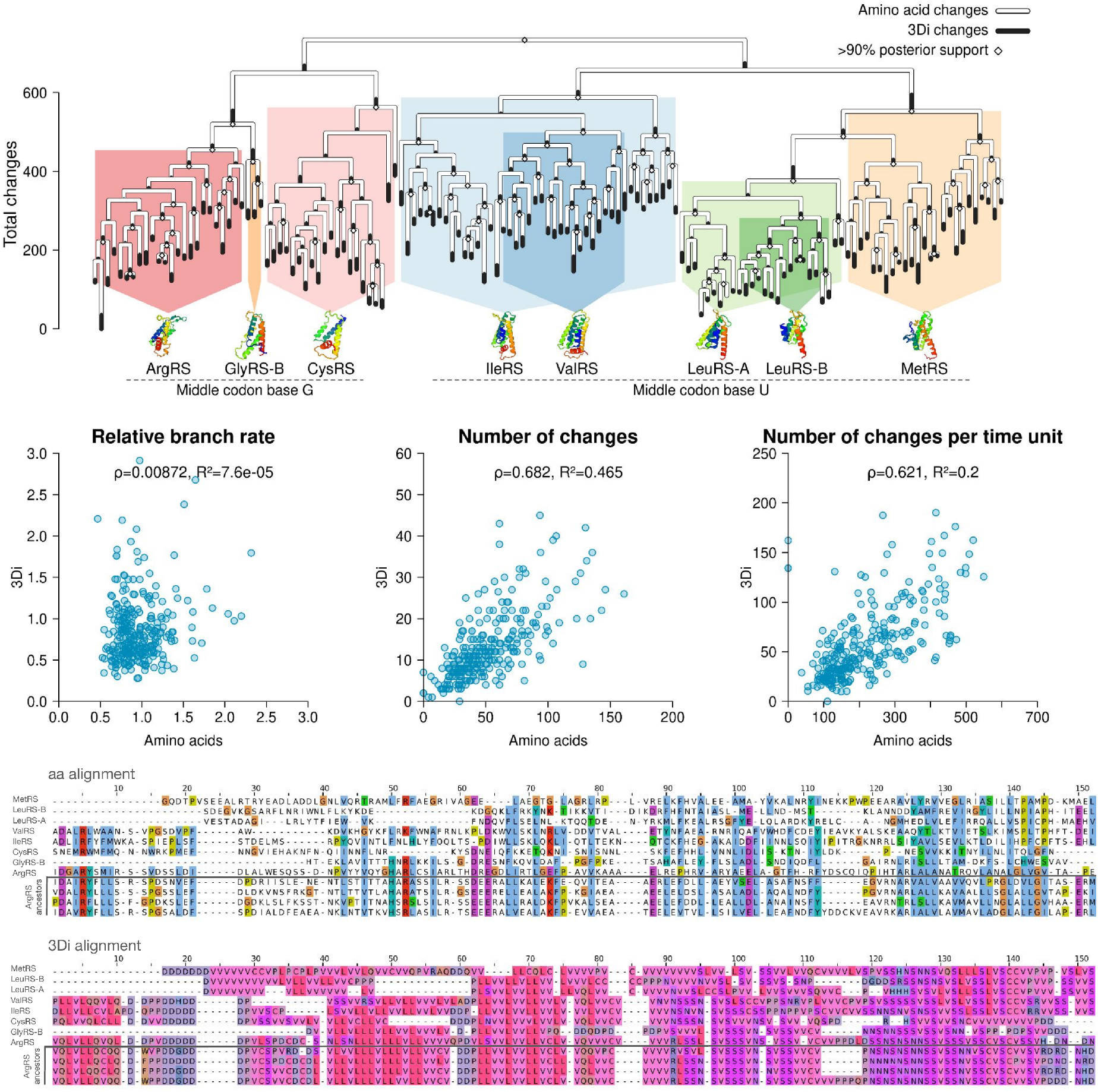
Top: Summary tree of the CRIMVLG anticodon binding domain. As expected, sequence changes (aa) are much more common than structural changes (3Di); reflecting the resilience of protein structure to mutation (Illergård et al., 2009). Middle row: We measured the per-branch correlation between aa and 3Di changes using three metrics, with the expectation to see a positive trend given the biochemical relationship between sequence and structure. First, we considered relative branch rates under the relaxed clock, which led to no association. Second, using BeastMap, we counted the number of changes on each branch, which led to a moderate association. Lastly, we divided the previous measure by branch length (in relative time units), which saw the same trend at a similar strength. In this instance, stochastic mapping provided a better estimate of “branch rate” than the branch-rate parameters of a relaxed clock model. Bottom: Snippets from the aa and 3Di alignments, with one full-length representative from each family. Shown are four ancestral sequence reconstructions for the ancestor of ArgRS, sampled from the posterior distribution. Alignments displayed with Jalview (Waterhouse et al., 2009).

These results also infer a different phylogeny of the CRIMVLG domain (Fig. 5) than the catalytic domain (Douglas et al., 2024a). This result may be due to phylogenetic uncertainty, or it may suggest a history of exchange of protein domains (Stolzer et al., 2015). Specifically, in this model the ValRS and LeuRS ABDs are, respectively, offshoots of the IleRS and MetRS clades. In contrast, their catalytic domains are each monophyletic. Interestingly, the ABDs associated with codon middle base G (ArgRS, GlyRS, and CysRS) form a distinct split in the tree from those with middle base U (IleRS, LeuRS, MetRS, ValRS), which may offer insights into how codons were assigned to amino acids during early code evolution (Kondratyeva et al., 2022).

### Case study 3: Spatial transmission of a respiratory virus

In 2022, New Zealand saw a surge of influenza cases following the reopening of the border (Huang et al., 2025), which was closed during the COVID-19 pandemic. One of these border incursions led to a large single-origin outbreak of influenza A(H3N2), represented by over 1300 fully sequenced genomes (Jelley et al., 2025). Due to the spatial nature of virus transmission, we would expect to observe a greater rate of transmission between areas that are close, compared with those that are further apart on the map. This is not only an intuitive expectation, but also has been well-characterised in New Zealand for other respiratory viruses too like SARS-CoV-2 (Geoghegan et al., 2020; Jelley et al., 2022) and respiratory syncytial virus (Jelley et al., 2024). This simple view of transmission does not take into account air travel, as earlier studies have done (Attwood et al., 2022).

To explore the spatial dynamics of influenza transmission, we downsampled the single-origin dataset from Jelley et al. (2025) into 96 sequences of the haemagglutinin segment (HA), each associated with a sample collection date and a Health Region (i.e., its *deme* / discrete location). Then, under a discrete trait biogeography model (Lemey et al., 2009), we estimated the rates of transmission among the four Health Regions – Northern (*N* ), Te Manawa Taki (*M* ), Central (*C*), and Te Waipounamu (*W* ). It is noted that this phylogeography model is quite sensitive to sample biases in comparison to structured coalescent or birth-death models (De Maio et al., 2015), but it will suffice for the purposes of this demonstration. In contrast to many previous approaches to this problem which associate transmission events only with nodes, by using stochastic mapping we can estimate the point along the lineage when these transitions occurred.

Using BeastMap, we estimated the points in time when these Health Region border crossing events occurred (Fig. 6). These results further corroborate the source-sink dynamic previously identified (Jelley et al., 2025), where the North Island regions (Northern, Manawa Taki, and Central) seeded transmission into the South Island (Te Waipounamu). Interestingly, the variation in migration count was much more pronounced than in migration rate. Among the four demes, there are 12 (asymmetric) possible migration event counts, which were estimated using stochastic mapping. The median estimates of these counts were 0 (*W* → *N, W* → *M* ), 1 (*W* → *C, M* → *W* ), 2 (*N* → *W, M* → *C*), 3 (*M* → *N* ), 4 (*C* → *N, C* → *W* ), and 5 (*N* → *C, N* → *M, C* → *M* ). By contrast, there are just 6 (symmetric) pairwise migration event rates, which are all estimated parameters; in units of migrations per lineage per year. The median estimates of these rates were 0.59 (*M* ↔ *W* ), 1.13 (*N* ↔ *W* ), 1.32 (*C* ↔ *W* ), 1.84 (*C* ↔ *M* ), 2.13 (*C* ↔ *N* ), and 2.32 (*N* ↔ *M* ). In addition to offering a more refined description (asymmetric as opposed to symmetric events), the counts were also more variable than the rates. The median counts have a coefficient of variation (i.e., variance divided by mean) of 1.4, while the rates are at just 0.13. The counts therefore appeared to be more sensitive to signal/noise (i.e., with a high variance) compared with the rates, likely due to their direct influence from the prior distribution.

**Fig. 6:**
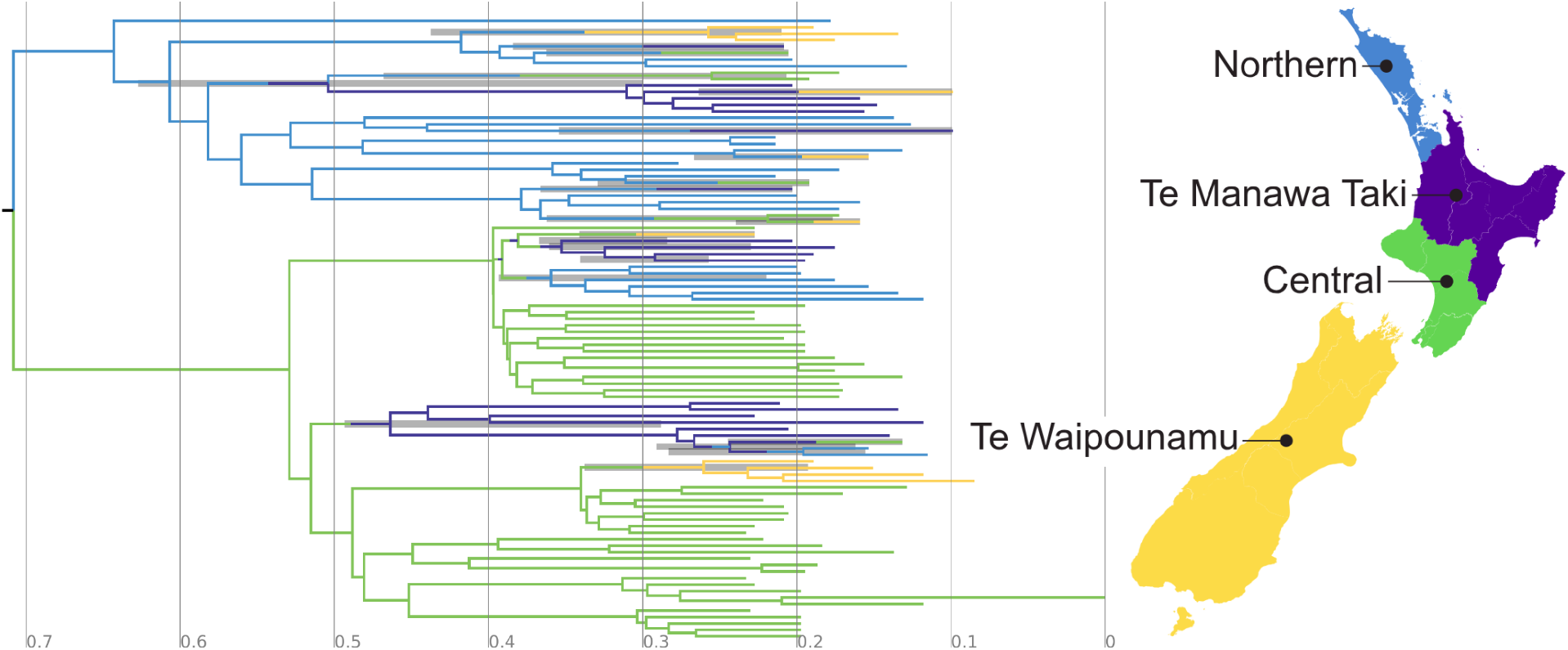
Summary of an influenza A(H3N3) outbreak in New Zealand, 2022, with lineages coloured by Health Region (shown on the nation’s map). The x-axis depicts the number of years since the most recent sample (24 Aug 2022). The posterior distribution of trees was summarised into a single point estimate using the CCD-0 method. Then, we summarised the posterior distribution of transitions between the four Health Regions onto the summary tree. The 95% credible interval of each border-crossing time is shown as a grey bar. Note that the moment of colour change on the branches is not a point estimate of when the transition occurred – rather the colour changes are conveniently positioned midway along the respective branch to prevent negative branch lengths. Only the 95% credible intervals depict time estimates. Tree picture generated by IcyTree (Vaughan, 2017).

The stochastic mapping approach reconstructs more influenza transmission between neighbouring demes than those more distant, in line with our expectation. The four demes here are arranged linearly on New Zealand’s map – starting at the north with N, through to M, C, and then W at the south. Now, let us define the *neighbouring* pairs of demes as: {*N, M*}, {*M, C*}, and {*C, W*}; and the *distant* pairs as: {*N, C*}, {*N, W*}, and {*M, W*}. We estimated a total of 20 transmission events between neighbours, and just 12 between distant locations (relative difference: 1.67). Whereas, the averaged rate of transmission was 1.83 between neighbours and 1.28 between distant areas (relative difference: 1.43). Both are consistent with our expectation that closer demes have more transmission than further ones. However, the discrepancy between neighbours and non-neighbours was more evident for counts than for rates. Further phylogenetic and data preparation details for this analysis can be found in Supporting Information.

## Discussion

Substitution histories – the evolutionary paths of sequence changes that have led to a set of observed sequences – are a core part of phylogenetic inference, whether branch lengths are expressed in terms of amount of change along branches (substitution trees), or in terms of absolute or relative time durations (time trees). In most phylogenetic methods, substitution histories and ancestral sequences are averaged over during inference and are obscured from the user output. However, in some cases we may be explicitly interested in reconstructing the history of changes, or the sequence states either at internal nodes of the tree or along the branches (Nielsen, 2002; Gumulya and Gillam, 2017; Attwood et al., 2022). But even where the history of changes is not the explicit focus, stochastic mapping provides an effective way of inferring amount of change, which is a core component of branch length estimation (Steel and Penny, 2000). Accurate branch length estimation is an essential part of phylogenetic inference not only in cases where absolute dates are required, such as origination of lineages or traits (Dos Reis et al., 2016) but also in cases where it is the relative timing of branching events that is of interest, such as diversification dynamics or phylogenetic diversity measures (Welch and Bromham, 2005; Duchêne et al., 2017; Ritchie et al., 2017).

We have implemented and validated a flexible software package, BeastMap, which can infer substitution histories within a Bayesian phylogenetic framework (i.e., during MCMC as opposed to a *post-hoc* analysis). Because it is implemented as a user-friendly addition to BEAST 2, it can be used in combination with a wide range of phylogenetic inference methods available in BEAST 2, including the multispecies coalescent (StarBeast3; Douglas et al. (2022)). Here we have demonstrated the application of the stochastic mapper within a number of different phylogenetic reconstruction methods, and validated them on simulated data that represent a range of evolutionary scenarios. Our trials indicate that substitution histories can be accurately and precisely estimated using BeastMap when estimating time trees using a relaxed clock (Fig. 2), compared with substitution trees, despite the various assumptions and requirements being violated by these datasets (Fig. 2 and Table S1–S4). However, in the case where sequences were simulated through unconstrained and un-clock-like processes, maximum likelihood substitution trees performed better than Bayesian gamma-length distributed substitution trees. Taken together, the parameter spaces explored in these simulation studies suggest that time trees using stochastic mapping are a useful way to infer branch lengths even when divergence dating is not the goal, notably under the optimised relaxed clock (ORC model), which has faster performance than other clock model implementations (Douglas et al., 2021).

In order to test the reliability of new methods, we need to apply them to cases where the true answer is known, so we can ask if the method can accurately recover the known history. We have used simulations under a range of models of sequence evolution, each of which departs from the assumptions of standard phylogenetic models in different ways. However, performance on simulated datasets does not necessarily reflect reliability on more complex real data. Here we road-test the stochastic mapper using three different empirical datasets that represent cases where we can evaluate the reasonableness of the inferred substitution histories on the basis of prior expectation. In the case of the mammalian sequences, we demonstrate that BeastMap reconstructs relatively more changes on branches leading to small bodied-lineages than their larger relatives, consistent with the widely observed phenomenon that smaller mammals tend to have a higher rates of molecular evolution than their larger relatives (Fig. 4; Bromham (2011); Cagan et al. (2022); Bergeron et al. (2023)). We also demonstrate the application of BeastMap to amino acid substitutions and structural change in a protein by studying the anticodon binding domains of aminoacyl-tRNA synthetases (Fig. 5). Our analysis shows that the rate of change in 3Di structural alphabet characters is associated with the amino acid substitution rate. This is an intuitive result, given the biological relationship between sequence and structure, and congruent with Moi et al. (2025), yet this correlation between sequence and structural change along each lineage was not apparent when comparing branch rates under the relaxed clock. BeastMap may therefore provide a useful tool for reconstructing deep history of protein evolution. Lastly, using the example of an influenza outbreak, we show how the stochastic mapper can be used to localise evolutionary transitions in space and time (Fig. 6). Stochastic mapping confirmed that this respiratory virus was more likely to spread to neighbouring Health Regions than distant ones, which is in line with common intuition around pathogen transmission. Other examples where stochastic mapping could be used to estimate ancestral sequences and substitution histories include cases of reconstructing steps in protein evolution (Laurin-Lemay and Rodrigue, 2025), inferring shifts in structure or function along phylogenies (Starr et al., 2017), or determining directional trends in sequence evolution (Jordan et al., 2005).

Estimating substitutions as counts along phylogenies has a number of potential advantages. Inferring branch-specific rates or lengths requires integrating observed data with prior distributions as part of the model, so they are dependent on model assumptions that in many cases are not informed by known evolutionary process but are convenient and empirically arbitrary models. In contrast, the number of substitutions occurring along a branch are counts, conditional on the inferred histories. In statistical inference, empirical quantities like counts are typically more sensitive to the data and consequently exhibit higher variance, whereas model-based estimates tend to be more conservative and may diminish any apparent trends. Consistent with this expectation, we found that substitution counts were generally more successful at recovering expected patterns in the cases studies than parametric rate estimates. For example, there was stronger evidence for a relationship between mammalian body size and substitution count, compared with parametric genetic distances. Likewise, a link between protein sequence and structural divergence was detectable using branch-wise substitution counts but not using relaxed-clock branch rates (Fig. 5). And, the faster spread of influenza to neighbouring Heath Regions over distant ones was more easily detected with stochastic mapping than with migration rates alone. At the same time, counts proved more difficult to estimate accurately than tree-wide parameters such as heights or total lengths (Table 1). Taken together, these case studies highlight how stochastic mapping can, in some cases at least, offer a more responsive view of evolutionary histories than rate-based summaries alone, revealing patterns that may be obscured or dampened when relying solely on parametric estimates.

The mammalian case study also illustrates another advantage of the stochastic mapping approach: an efficient and user-friendly way to estimate the number synonymous and non-synonymous changes along branches of a phylogeny from alignments of protein coding sequences. This approach is computationally efficient because it avoids the need to estimate *ω* directly through a computationally demanding 61 × 61 matrix (Goldman and Yang, 1994). Surprisingly, we found that estimating branch lengths (in substitutions per site) were much less challenging than the actual number of substitutions via stochastic mapping, meaning that the latter made a much more stringent test for benchmarking phylogenetic methods (Table 1). Within the set of tests performed here, analysing protein-coding genes as nucleotides (i.e., with 4 states) was a reliable alternative to a codon-based model (i.e., with 61 states) when using codon partition models (Shapiro et al., 2006) in conjunction with free rates (Soubrier et al., 2012). This further corroborates the observation that gamma heterogeneity gives an over-simplistic description of site rate variability (Ferretti et al., 2024). Our estimator of 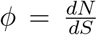, which is simply counted as 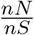 times a constant term, offers a reliable estimation of *ω* in just a fraction of the runtime (from 15 to 40 times faster in our mammalian datasets). Despite its simplicity, *ϕ* also gives a low-bias estimate of *ω* (Fig. S2), and even outperformed some of the more elegant approaches described by Lemey et al. (2012) that are also based on nucleotide stochastic mapping. Because the rate of synonymous substitutions is expected to be primarily driven by the mutation rate (while non-synonymous rates are influenced by mutation rate, population size and selection), finding a consistent association between body size and synonymous substitution rate confirms that mutation rates in mammals scale with life-history.

The case study with aminoacyl-tRNA synthetases provides a different example of estimating and comparing number of changes in different classes, in this case amino acid substitutions and structural changes. This could be used to examine the relative rates of change of sequence and structural change in proteins, but it could also be applied more generally to examining the influence of different factors on the rate of structural changes. Given that structural changes provide one potential solution to the challenges of reconstructing evolution in deep time (Puente-Lelievre et al., 2025), reconstructing substitution histories of inferred structural changes provides a way of examining the rate of change over time.

One major limitation in stochastic mapping comes from indels. The important biological process of insertion and deletion is generally overlooked in phylogenetics (Redelings et al., 2024). Indels are usually ignored and treated as missing data (i.e., clades with all-gaps at a given site are omitted from the site likelihood calculation). While this might not always influence phylogenetic inference (Truszkowski and Goldman, 2016; Machado et al., 2021), it can be quite problematic for ancestral sequence reconstruction and stochastic mapping. This is a major challenge for those who seek to study gene functions in a laboratory environment; as how one characterises indels directly translates to the size of a gene, which will impact protein folding and function. Consequently, there are programs for performing ASR with indel models built in, such as FastML (Ashkenazy et al., 2012) and ARPIP (Jowkar et al., 2023). If deletions were to be treated as ambiguous characters during stochastic mapping (e.g., the ‘N’ nucleotide), the ancestral sequence lengths will be overestimated (because deletions are being ignored), and so too will the number of substitutions along each lineage. One solution is to remove all gapped sites and/or incomplete sequences from the alignment (as we did for our mammalian datasets). However, this can result in large volumes of data being discarded. But if these gaps indeed represent historical insertion and deletion events, and are not just missing data, then a more elegant solution is through the use of a phylogenetic indel model (as we did for our anticodon binding domain analysis). However, this necessitates even further modelling decisions to be made, such as the choice of indel model and its various prior distributions; which may in turn have major impacts on the phylogeny. Our implementation represents indels using the simple indel coding described by Simmons and Ochoterena (2000), however alternative models may be more reasonable, such as the complex indel coding method (Simmons and Ochoterena, 2000) or Poisson indel processes (Jowkar et al., 2023). The handling of gaps remains a major challenge in stochastic mapping.

In summary, we have provided a new user-friendly implementation of stochastic mapping and demonstrated how the framework can offer new insights into evolutionary processes, complementary to what we can learn from substitution rates. We have shown that relaxed clocks provide a powerful means for estimating those histories, even when temporal calibration and divergence dating is not part of the objective. Future innovations may see the continued development of explicit substitution models (Varilly et al., 2025), which will enable the estimation of substitution histories without the need for stochastic mapping. With the continued development of molecular clock, indel evolution, and tree branching models and model-comparison frameworks, it is hoped that substitution history reconstruction will become a routine part of phylogenetic analysis.

### Software and Data Availability

Our software package is implemented as a package within BEAST 2, and can be configured through a user-friendly graphical interface via BEAUti. This enables seamless integration with a range of existing data types, site, clock, and tree models during MCMC available in BEAST 2 (Bouckaert et al., 2019). The graphical interface also supports an indel model whereby gaps are represented using the Simple Indel Coding method (Simmons and Ochoterena, 2000), which can overcome estimation biases that come with treating gaps as ambiguous characters (the default). In addition to performing ASR, BeastMap can also report the total sum of changes along each lineage, their times, and the sum of different kinds of change, such as synonymous and non-synonymous, transition and transversion; while offering further user-specified options such as counting the number of substitutions from a state in set *x* to *y* (e.g., from an acidic amino acid *x* = {D, E} to a basic one *y* = {H, K, R}). A counter can also be restricted to consider only specific sets of positions in a sequence. The method is compatible with a wide range of discrete data types, including nucleotide, codon, amino acid, 3Di, morphological, cognate, phoneme, and indel. Graphical user interface support (BEAUti) is provided for standard BEAST 2 analyses, as well as the StarBeast3 multispecies coalescent (Douglas et al., 2022).

Code and documentation for BeastMap is available at http://github.com/jordandouglas/BeastMap. The 3Di substitution model averaging framework, which compares empirical substitution matrices, is available at http://github.com/jordandouglas/FoldBeast. Datasets are available on Dryad.

## Acknowledgements

The authors thank Remco Bouckaert and Xia Hua for their helpful feedback; and Lauren Jelley, PHF Science, and the New Zealand Ministry of Health for providing the influenza virus data.

## Supporting Information

Further methodological details, including prior distributions, are provided in the Supporting Information.

