## Supporting Information for "Reconstructing substitution histories on phylogenies, with accuracy, precision, and coverage"

December 22, 2025

#### Contents

**1 Gamma-length tree prior** **3**

**2 Benchmark results** **6**

**3 dN/dS estimators** **9**

**4 Phylogenetic models and their prior distributions** **10**

### 1 Gamma-length tree prior

Let $\mathcal{T}$ be a binary rooted tree with $n$ leaves and $n-1$ internal nodes. These nodes $X = (x_1, \ldots, x_{2n-1})$ are numbered starting from the leaves at $x_1, \ldots, x_n$, followed by internal nodes, and finally the two children of the root and the root itself $x_{2n-3}, x_{2n-2}, x_{2n-1}$. Let $L_{x_i} > 0$ be the length of the branch between $x_i$ its parent, where $L_{x_{2n-1}} = 0$ at the root. Suppose here that these branch lengths are in units of substitutions per site and hence $\mathcal{T}$ is a *substitution tree*.

Under the gamma-length tree prior, branch lengths are independent and identically distributed from a gamma distribution. The tree is rooted following inspiration from the midpoint rooting method. First, we will describe a generative process for directly sampling a rooted gamma-length tree from a distribution. Then, we will provide the probability density function. Lastly, we will describe proposal kernels for estimating $\mathcal{T}$ using the Metropolis-Hastings MCMC algorithm.

The construction of a gamma-length midpoint tree on $n$ taxa is much like a pure birth tree:

1. Let $S \leftarrow \{x_1, x_2, \ldots, x_n\}$ be our set of subtrees (initially just leaves)
2. Let $i \leftarrow n + 1$ be our next node index
3. While $|S| > 1$:
   (a) Sample $a, b \in S$ uniformly at random, without replacement
   (b) $c \leftarrow x_i$
   (c) If $|S| > 2$:
      i. Add an edge between $a$ and $c$ with length $L_a \sim \text{Gamma}(\text{shape} = \kappa, \text{scale} = \beta)$
      ii. Add an edge between $b$ and $c$ with length $L_b \sim \text{Gamma}(\text{shape} = \kappa, \text{scale} = \beta)$
   (d) Else:
      i. Sample the sum of root edges $L \sim \text{Gamma}(\text{shape} = \kappa, \text{scale} = \beta)$
      ii. Add an edge between $a$ and the root $c$ with length $L/2$
      iii. Add an edge between $b$ and the root $c$ with length $L/2$
   (e) $S \leftarrow S \setminus \{a\}$
   (f) $S \leftarrow S \setminus \{b\}$
   (g) $S \leftarrow S \cup \{c\}$
   (h) $i \leftarrow i + 1$
4. Let $Y = (y_1, y_2, \ldots, y_k)$ be the path between the furthest pair of leaves in the tree rooted at $S$
5. Let $D \leftarrow \sum_j L_{y_j}$ be the length of this path
6. Let $F \sim \text{Beta}(\text{shape1} = \alpha, \text{shape2} = \alpha)$
7. Reroot the tree at length $FD$ along the path from $y_1$ to $y_k$

In this construction, branch lengths are independent and identically drawn from a gamma distribution with shape $\kappa$ and a mean of $\kappa\beta$. The root of the tree is approximately halfway along the longest path between any two taxa, or more precisely $0 < F < 1$ of the way, where $F \sim$ Beta$(\alpha, \alpha)$ for some $\alpha \geq 1$. When $\alpha = 1$, the root can lie anywhere along the longest path, with equal probability. As $\alpha$ approaches infinity, the tree approaches a perfectly equidistant midpoint tree. For our purposes, let us say that $\alpha = 50$, meaning that $F$ has a 95% interquartile range of (0.4, 0.6). It is noted that in a typical phylogenetic model, with a time-reversible substitution model and no further prior information on tree topology, all of our information on root placement will come from the user-defined $\alpha$, which we will fix.

The probability density of $\mathcal{T}$, including its branch lengths $L$ and relative root placement $F$, is:

$$p\Big(\mathcal{T} \,\Big|\, \alpha, \kappa, \beta\Big) = p\Big(F \,\Big|\, \alpha\Big) \times \sum_{i=1}^{2n-4} p\Big(L_{x_i} \,\Big|\, \kappa, \beta\Big) \times p\Big(L_{x_{2n-3}} + L_{x_{2n-2}} \,\Big|\, \kappa, \beta\Big), \tag{1}$$

where $p(F|\alpha)$ is the probability density function of a Beta($\alpha$, $\alpha$) distribution, and $p(L|\kappa, \beta)$ is the density of a Gamma($\kappa$, $\beta$).

Finally, generating proposals on $\mathcal{T}$ can be performed using the operations below. Where applicable, these operators make use of the Bactrian proposal kernel for selecting step sizes, which has been shown to be around 15% more efficient at achieving convergence compared with than normal or uniform distributions.[1]

1. `TipRandomWalker`: alter the length of a randomly selected leaf by $w$, which is derived from a Bactrian distribution.

2. `ScaleLengths`: multiply a randomly selected branch by a scale factor $s$, which is derived from a Bactrian distribution.

3. `ScaleHeights`: multiply all node heights (relative to the youngest leaf) by a constant value $s$ (Bactrian). This is based on a standard operator in BEAST 2,[2] but with the Hastings ratio adjusted to account for the fact that our probability density is on branch lengths not node heights.

4. `AVMN`: Similar to `ScaleHeights`, except using an adaptive method (ref) that approximates the correlations between each parameter (in this case $2n-1$ node heights, $\kappa$, and $\beta$) using a multivariate normal distribution.

5. `EpochFlex` and `EpochStretch`: highly efficient tree length operators introduced in the BICEPS package for BEAST 2.[3]

6. `NarrowExchange`, `WideExchange`, `SubtreeSlide`, `WilsonBalding`, and `BactrianNodeOperator`: standard tree topology and branch lamsmath amssymbength operators in BEAST 2.

As shown in the main results of this paper, and Fig. S1, it turns out that this gamma-length tree prior is not suitable for describing clock-like evolution

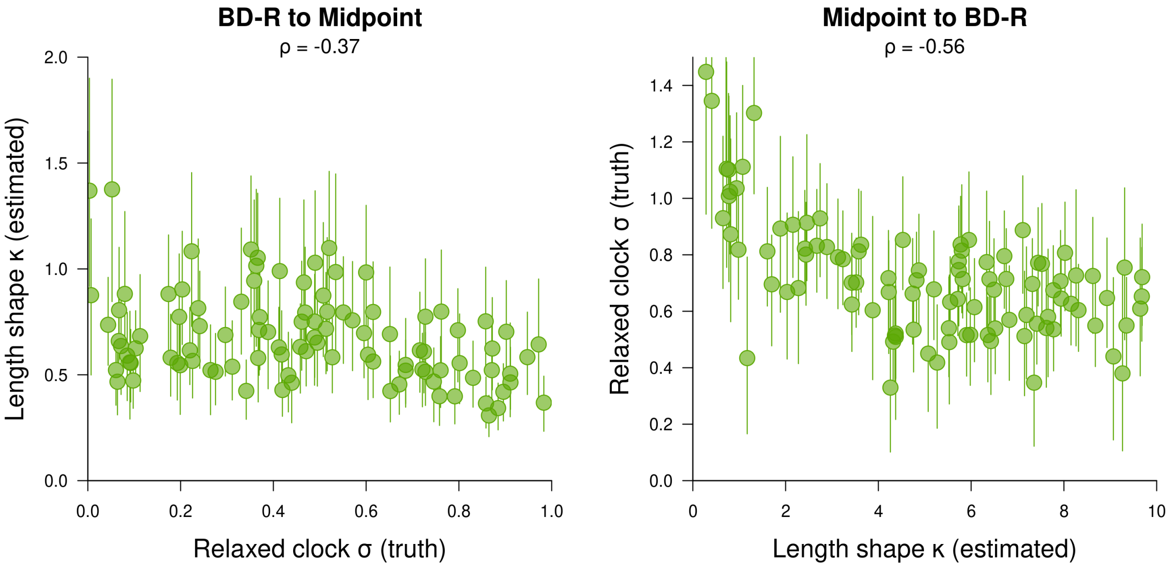

Fig. S1: There is not a well-defined relationship between $\kappa$ (the branch length shape from the gamma-length prior) and $\sigma$ (the branch rate standard deviation under a relaxed clock). Each point represents a single MCMC chain performed on a dataset simulated under **BD-R** or **Midpoint**, with inference done under the opposite; with circles depicting median estimates and verivtal lines as 95% credible intervals. Although there is a clear negative relationship between $\sigma$ and $\kappa$, this relationship is inconsistent. When the true $\sigma$ is close to 0 (corresponding to a strict clock), the estimated $\kappa$ is close to 1. But when the true $\kappa$ is close to 1, the estimated $\sigma$ is close to 0.8 (highly variable rates).

#### 2 Benchmark results

| Simulator | Substitutions | Estimator | N. obs | Coverage (%) | Correlation | Mean Rel. Error | SD Rel. Error |
|---|---|---|---|---|---|---|---|
| BD-R | Total | ML | 100 | 26 | 0.964 | 7.19 | 0.0598 |
| | | GM | 100 | **96** | 0.732 | 5.36 | 0.0489 |
| | | BD-S | 100 | 85 | 0.330 | 5.74 | 0.0427 |
| | | BD-R | 100 | **95** | **0.985** | **4.15** | **0.0339** |
| | Leaves | ML | 4000 | 69.2 | 0.908 | 23.7 | 0.28 |
| | | GM | 4000 | **95** | 0.829 | 23.3 | 0.257 |
| | | BD-S | 4000 | 85 | 0.368 | 24.6 | 0.301 |
| | | BD-R | 4000 | **94** | **0.950** | **19.6** | **0.212** |
| | Internal | ML | 2896 | 56.6 | 0.914 | 46.2 | 0.617 |
| | | GM | 2717 | 88.5 | 0.828 | **38.5** | **0.482** |
| | | BD-S | 2977 | 75 | 0.310 | 48.4 | 0.697 |
| | | BD-R | 2999 | 86.5 | **0.932** | 39.5 | 0.533 |

Table S1: **BD-R** simulator benchmarking. Substitution count estimators were benchmarked on 400 simulated datasets. The best score in each band is highlighted in bold, with the exception of Coverage, which is bold when within the range of 91–99%. The number of observations is 100 for the full tree (one per dataset/tree), 4000 for leaves (40 leaves per tree and 100 trees), and variable for the internal nodes (as the estimates of substitution counts on internal nodes are conditional on the clade having over 50% support). All statistics are rounded to three significant figures (except for N. obs).

| Simulator | Substitutions | Estimator | N. obs | Coverage (%) | Correlation | Mean Rel. Error | SD Rel. Error |
|---|---|---|---|---|---|---|---|
| BD-C | Total | ML | 100 | 43 | 0.906 | 8.72 | 0.151 |
| | | GM | 100 | 90 | 0.866 | 5.15 | 0.12 |
| | | BD-S | 100 | 87 | **0.931** | 4.63 | **0.0417** |
| | | BD-R | 100 | 89 | 0.93 | **3.97** | 0.0426 |
| | Leaves | ML | 4000 | 80.4 | 0.923 | 16 | 0.227 |
| | | GM | 4000 | **96** | 0.962 | 19 | 0.26 |
| | | BD-S | 4000 | **92.8** | 0.959 | 16.4 | 0.192 |
| | | BD-R | 4000 | **95.8** | **0.97** | **15.3** | **0.19** |
| | Internal | ML | 2989 | 68.2 | 0.916 | 42.2 | 0.632 |
| | | GM | 2987 | **91.6** | **0.932** | **30.6** | **0.408** |
| | | BD-S | 3009 | 83.8 | 0.914 | 46.4 | 0.774 |
| | | BD-R | 2728 | 89.2 | 0.929 | 38.4 | 0.6 |

Table S2: **BD-C** simulator benchmarking. Refer to Table S1 caption for notation.

| Simulator | Substitutions | Estimator | N. obs | Coverage (%) | Correlation | Mean Rel. Error | SD Rel. Error |
|---|---|---|---|---|---|---|---|
| ME-CG | Total | ML | 100 | 23 | 0.963 | 6.95 | 0.0434 |
| | | GM | 100 | 76 | 0.137 | 10.1 | 0.246 |
| | | BD-S | 100 | 79 | 0.822 | **5.51** | **0.0348** |
| | | BD-R | 100 | 78 | **0.980** | 5.86 | 0.0362 |
| | Leaves | ML | 4000 | 61.1 | 0.901 | 17 | 0.121 |
| | | GM | 4000 | 85.7 | 0.130 | 19.4 | 0.213 |
| | | BD-S | 4000 | 75.3 | 0.833 | 15.1 | 0.0961 |
| | | BD-R | 4000 | 87.8 | **0.940** | **14.3** | **0.0931** |
| | Internal | ML | 2412 | 35.9 | 0.741 | 139 | 1.93 |
| | | GM | 2143 | 59.4 | 0.0561 | 123 | 1.71 |
| | | BD-S | 2277 | 48.6 | 0.743 | 129 | 1.68 |
| | | BD-R | 2234 | 54.7 | **0.783** | **116** | **1.47** |

Table S3: **ME-CG** simulator benchmarking. Refer to Table S1 caption for notation.

| Simulator | Substitutions | Estimator | N. obs | Coverage (%) | Correlation | Mean Rel. Error | SD Rel. Error |
|---|---|---|---|---|---|---|---|
| GM | Total | ML | 100 | 52 | **0.991** | **3.71** | 0.0531 |
| | | Midpoint | 100 | 75 | 0.681 | 5.9 | 0.0831 |
| | | BD-S | 100 | 90 | 0.601 | 4.4 | **0.043** |
| | | BD-R | 100 | **94** | 0.935 | 4.78 | 0.155 |
| | Leaves | ML | 4000 | 76.8 | **0.909** | **16.5** | **0.181** |
| | | Midpoint | 4000 | **91.4** | 0.742 | 18.9 | 0.195 |
| | | BD-S | 4000 | 77.3 | 0.516 | 28.3 | 0.371 |
| | | BD-R | 4000 | **93** | 0.883 | 17.2 | 0.19 |
| | Internal | ML | 3347 | 73.7 | **0.921** | 24 | **0.255** |
| | | GM | 3269 | 84.8 | 0.717 | 27.7 | 0.315 |
| | | BD-S | 3040 | 77.3 | 0.486 | 30.8 | 0.327 |
| | | BD-R | 3253 | **92** | **0.921** | **23.4** | 0.258 |

Table S4: **GM** simulator benchmarking. Refer to Table S1 caption for notation.

### 3 dN/dS estimators

We compared several $\frac{dN}{dS}$ estimators, as outlined in the Results section of the main article. These results are presented in Fig. S2, and suggest that $\phi$ makes a useful estimator of $\omega$.

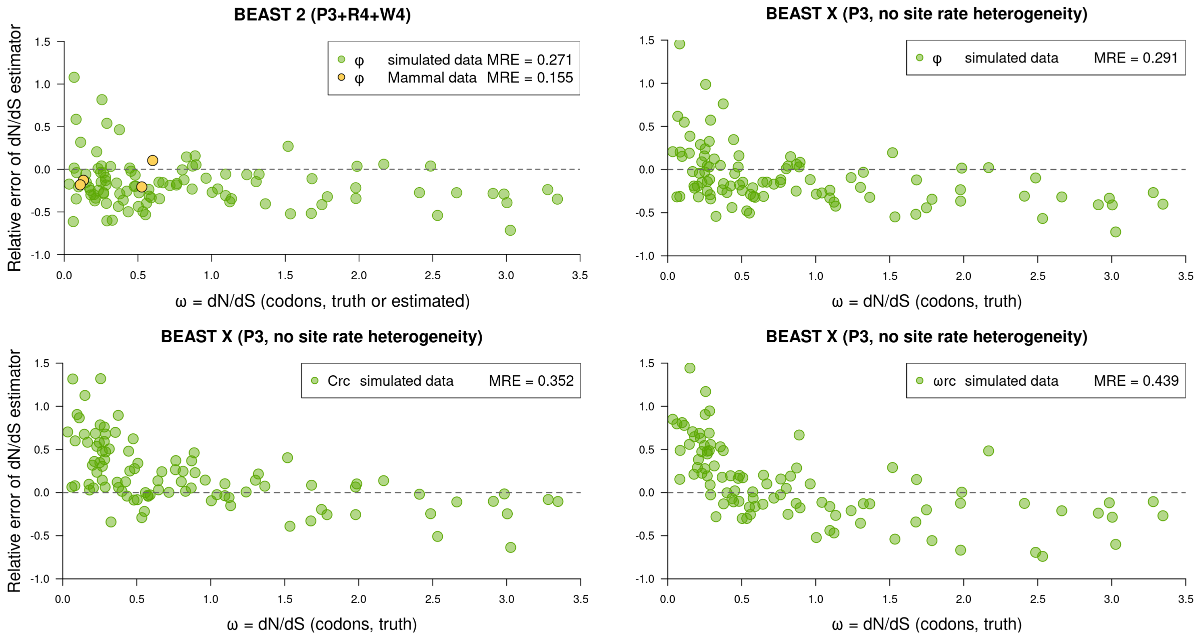

Fig. S2: Comparison of $\frac{dN}{dS}$ estimators based on nucleotide sequences. In the top two panels, we benchmarked our introduced $\phi$ term against values of $\omega$ that are either known (simulated codon data) or estimated (from Mammalian codon data). This term was calculated using the stochastic mappers in BEAST 2 and BEAST X. The mean relative error (MRE) is reported. We compared this estimator with the $C^{\mathrm{RC}}$ and $\omega^{\mathrm{RC}}$ estimators described by.[4] Those two estimators (without site rate heterogeneity) are implemented in BEAST X, but not BEAST 2. Collectively, these results further corroborate that the P3+R4+W4 model gives a better approximation of $\frac{dN}{dS}$ than P3 without heterogeneity. This experiment also confirms that $\phi$ appears to give a better estimate of $\omega$ than $C^{\mathrm{RC}}$, which is in turn more accurate than taking the site-wise average of $\omega^{\mathrm{RC}}$.

### 4 Phylogenetic models and their prior distributions

#### 4.1 Well-calibrated simulation studies

For our coverage studies (Fig. 3 of main article), the evolutionary model was composed of a gamma-length / midpoint tree prior, a strict clock, an HKY substitution model with estimated frequencies, and four gamma site rate heterogeneity categories. We validated this approach using coverage simulation studies with 30 taxa and 300 sites. As shown in Fig. S3, these experiments confirmed the true value of a parameter lies in its 95% credible interval approximately 95% of the time; thereby providing confidence in the correctness of our method, and its ability to recover parameter estimates from data simulated under a known model.

During these experiments, the following priors were used:

- Branch shape $\kappa \sim \text{LogNormal}(\text{mean} = 0.05, \sigma = 1)$
- Branch length mean $\kappa\beta \sim \text{LogNormal}(\text{mean} = 2, \sigma = 1)$
- Gamma site rate heterogeneity shape $\sim \text{Exponential}(\mu = 1)$
- HKY transition-transversion ratio $\sim \text{LogNormal}(\text{mean} = 3, \sigma = 0.5)$
- Nucleotide equilibrium frequencies $\sim \text{Dirichlet}(\alpha_A = 2, \alpha_C = 2, \alpha_G = 2, \alpha_T = 2)$

Under these conditions, the mean tree height was 0.45 substitutions per site, with a 95% HPD (highest posterior density) interval of (0.023, 1.47). The midpoint prior $\alpha$ was fixed to 50, meaning that the root *a priori* resided between 40 – 60% along the path between the two furthest leaves (with 95% probability).

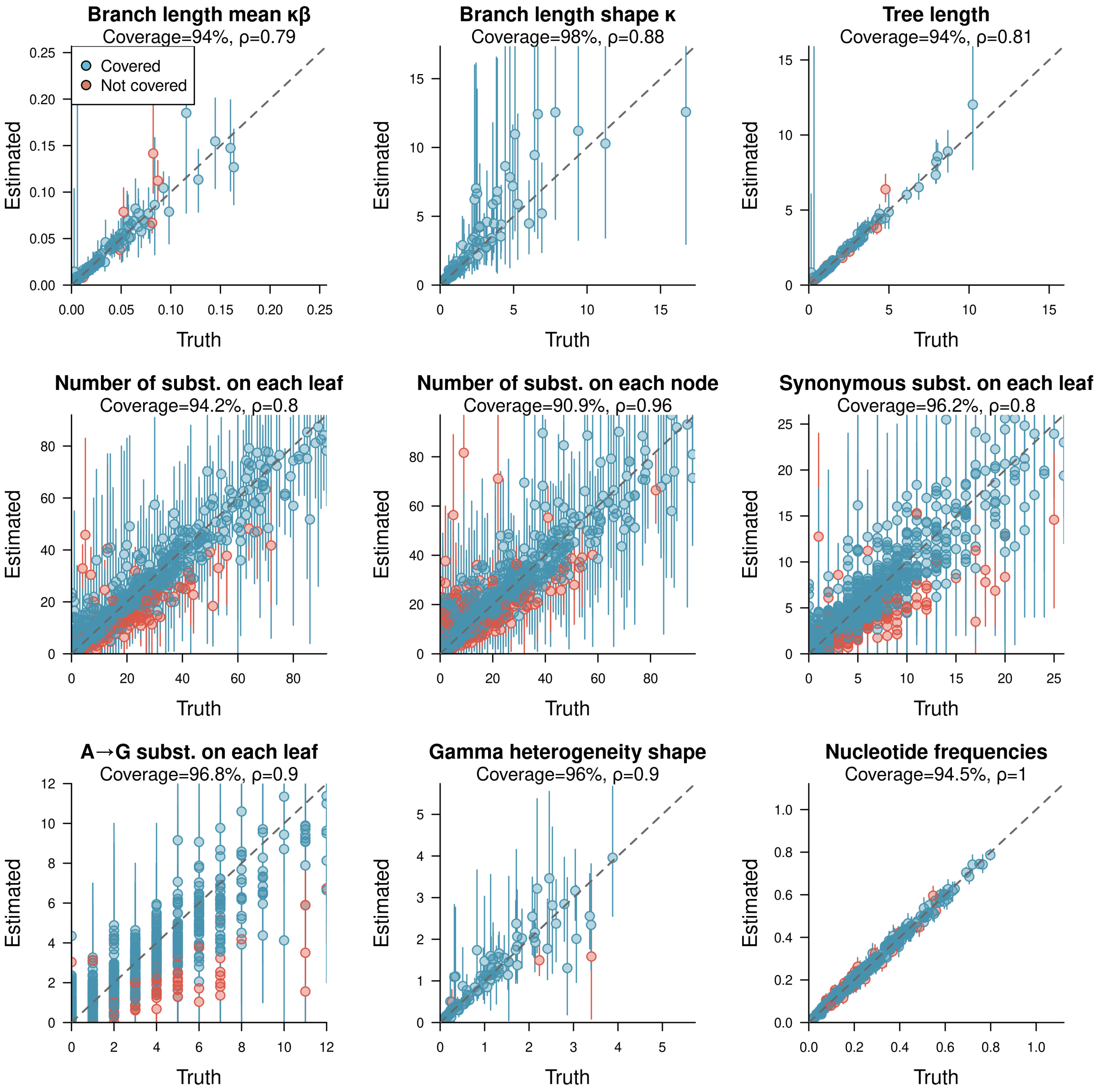

Fig. S3: Validating the gamma-length tree prior and stochastic mapper implementation. Each point represents a single MCMC chain run on one of 100 simulated datasets each with 30 taxa and 300 nucleotide sites. Mean estimates are shown by circles and 95% credible intervals by vertical lines. The coverage is the percentage of replicates where the true value is within the 95% credible interval, and $\rho$ coefficient measures the Pearson correlation between true and mean values. Some metrics are combined into a single panel – the substitution (subst.) counts are combined across each leaves or internal node, and frequencies are combined across the four characters. In the case of internal nodes (first row, second column), each point corresponds to a single clade in a true tree that appeared at least 30 times in a posterior distribution of trees.

#### 4.2 Performance benchmarking datasets

All benchmarking datasets consisted of 40 taxa and 300 nucleotide sites. The site model consisted of an HKY substitution model with frequencies estimated, and four gamma site rate categories (with the same prior distributions that were used for the well-calibrated simulation studies). 100 datasets were simulated under each of the conditions below, coming to a total of 400 datasets. In each case, the model (including its priors) used during simulation differs from those used during inference.

##### 4.2.1 BD-R datasets

The **BD-R** datasets were simulated from time trees, and with sequences that range from very clock-like to very non-clock-like. The time trees were simulated using a birth-death (**BD**) model,[5] and their sequences from a relaxed (**R**) clock. The birth-death model assumes that new species are born with rate $\lambda$ and die with rate $\mu$, where both terms are constant through time. These parameters were simulated from $\lambda \sim \text{Log-normal}(2.18, 0.5)$ and $\frac{\lambda}{\mu} \sim \text{Uniform}(0.8, 5.0)$ distributions at the start of each simulation. Under these conditions, the average tree height was 0.65 substitutions per site, with a 95% HPD of (0.1, 1.6). Under the uncorrelated relaxed clock,[6,7] lineages were assigned relative clock rates independently and identically distributed from a log-normal distribution, with a mean-in-real-space of 1 and a standard deviation $\sigma$, where $\sigma \sim \text{Uniform}(0, 1)$. As $\sigma$ approaches 0, the rate variation becomes smaller and the sequences become more clock-like. In practice, this is often achieved when $\sigma \lesssim 0.3$. When $\sigma \gtrsim 0.8$, the sequence violate the molecular clock hypothesis rather strongly. We previously estimated $\sigma$ ranging from 0.19 to 0.53 across eight empirical datasets.[7]

##### 4.2.2 BD-C datasets

The **BD-R** datasets were also simulated from birth-death time trees, and with a local (correlated) clock model.[8] Trees were simulated from $\lambda \sim \text{Log-normal}(\text{mean} = 20, \sigma 0.5)$ and $\frac{\lambda}{\mu} \sim \text{Uniform}(0.8, 5.0)$ distributions at the start of each simulation. Under these conditions, the average tree height was 0.32 substitutions per site, with a 95% HPD of (0.07, 0.82). The average clock rate was 1 substitution per site per unit of time across the tree, however the clock rate would differ slightly between each node and its parent. The clock rate of each branch is equal to its parent's rate times $M$, where $M$ is log-normally distributed with a mean-in-real-space of 1 and a standard deviation of 0.3.

##### 4.2.3 GM datasets

The **GM** datasets are potentially quite evolutionarily unrealistic. They are not generated through any explanatory temporal process, and thus serve as an extreme case for how the estimators will behave when so many standard assumptions of evolution are being violated. These substitution trees were simulated under the gamma-length / midpoint prior. Branch lengths were independent and identically distributed under a gamma distribution, with shape $\kappa \sim \text{Uniform}(0, 100)$ and mean length $\beta\kappa \sim \text{Log-normal}(-2.9, 1.0)$ at the start of each simulation. The midpoint rooting $\alpha$ was fixed at 50. Under these conditions, the average tree height was 0.49 substitutions per site, with a 95% HPD of (0.02, 1.9). To ensure that the branches describe expected number of substitutions we applied a strict clock model.

###### 4.2.4 ME-CG datasets

The **ME-CG** datasets make some pathological departures from standard phylogenetic assumptions, while maintaining some level of biological realism. First, these time trees were simulated under the assumption of two mass extinction (**ME**) events and three epochs of birth-death rates $\lambda$ and $\mu$. Second, the sequences were simulated along a clock gradient (**CG**) where the clock rate increases or decreases through time on average, in a noisy fashion. Trees were simulated using the TESS package for R.[9] Within this framework, birth-death trees were simulated with a maximum duration of $T = 0.5$ time units. Then, two mass extinction events times $t_1, t_2$ were sampled independently from $\frac{t_1}{T}, \frac{t_1}{T} \sim$ Beta$(2, 2)$, and two epoch change times $t_3, t_4$ from $\frac{t_3}{T}, \frac{t_4}{T} \sim$ Beta$(2, 2)$. At extinction event $i$, the proportion of lineages that survive are $p_i \sim$ Beta$(2, 2)$. The birth and death rates across the three epochs are $\lambda = (1, 3, 2)$ and $\mu = (0.9, 1, 2.1)$, going forward in time. Lastly, the proportion of extant taxa that are included in our sample of size $n = 40$ (i.e., rho-sampling) is also drawn from a Beta$(2, 2)$. Sequences were simulated following a local clock.[8] The average clock rate was 1 substitution per site per unit of time across the tree, however the average rate would either increase or decrease through time according to a gradient $G$. The clock rate of each branch is equal to its parent's rate (with 0.5 probability), or its parent's rate times $M$ (with 0.5 probability), where $M$ is log-normally distributed with a mean-in-real-space of $G$ and a standard deviation of 0.2. $G$ is also log-normally distributed with a mean-in-real-space of 1 and a standard deviation of $0.1$. Thus, when $G > 1$, the clock rate increases through time on average, and when $G < 1$, it decreases.

##### 4.3 Performance benchmarking models

Each model below is a misspecification of the simulated datasets they are benchmarked on. In all cases, an HKY substitution model with estimated frequencies and four gamma site rate categories were used. In the case of the Bayesian methods, the following priors were used for the site model:

- Gamma site rate heterogeneity shape $\sim$ Exponential$(\mu = 1)$
- HKY transition-transversion ratio $\sim$ LogNormal$(\text{mean} = 3, \sigma = 0.5)$
- Nucleotide equilibrium frequencies $\sim$ Dirichlet$(\alpha_A = 2, \alpha_C = 2, \alpha_G = 2, \alpha_T = 2)$

###### 4.3.1 ML model

The maximum likelihood substitution trees were found using IQ-tree.[10] As this is a maximum likelihood tree, there are no prior distributions. The site model was fixed to `HKY+G4+F` with the `-gmedian` option enabled to ensure compatibility with the BEAST 2 implementation of gamma site rate heterogeneity.

###### 4.3.2 GM model

In the gamma-length midpoint tree prior, the following prior distributions were used. A strict clock was applied here, with a clock rate fixed at 1.

- Branch shape $\kappa \sim$ LogNormal$(\text{mean} = 0.05, \sigma = 1)$
- Branch length mean $\kappa\beta \sim$ LogNormal$(\text{mean} = 2, \sigma = 1)$

###### 4.3.3 BD-S model

In the birth-death strict clock model, the following prior distributions were used. The clock rate was fixed at 1.

- Speciation rate $\lambda \sim$ LogNormal$(\text{mean} = 1, \sigma = 2)$
- Reproduction number $\frac{\lambda}{\mu} \sim$ LogNormal$(\text{mean} = 3, \sigma = 0.5)$

###### 4.3.4 BD-R model

In the birth-death (uncorrelated) relaxed clock model, the following prior distributions were used. The clock rate was fixed at 1.

- Speciation rate $\lambda \sim$ LogNormal$(\text{mean} = 1, \sigma = 2)$
- Reproduction number $\frac{\lambda}{\mu} \sim$ LogNormal$(\text{mean} = 3, \sigma = 0.5)$
- Branch rate standard deviation $\sigma \sim$ LogNormal$(\text{mean} = 0.4, \sigma = 0.5)$

##### 4.4 nN/nS simulation studies

The 100 codon datasets were simulated under a codon substitution model with four gamma rate categories (M0+F61+G4[11]), with each codon position following a general time reversible (GTR) substitution model.[12] Each dataset consisted of 200 codons (and 600 nucleotides) across 30 taxa. Trees were simulated under a birth-death tree prior and a relaxed clock (with a branch rate standard deviation of 0.2).

During simulation (codon-based), the following priors were used:

- Speciation rate $\lambda \sim \mathrm{LogNormal}(\mathrm{mean} = 10, \sigma = 0.5)$
- Reproduction number $\frac{\lambda}{\mu} \sim \mathrm{LogNormal}(\mathrm{mean} = 4, \sigma = 0.5)$
- $\omega \sim \mathrm{LogNormal}(\mathrm{mean} = 1, \sigma = 1)$
- Gamma site rate heterogeneity shape $\sim$ Exponential$(\mu = 1)$
- GTR relative rates $\sim$ Dirichlet$(\alpha_{AC} = 1, \alpha_{AG} = 4, \alpha_{AT} = 1, \alpha_{CG} = 1, \alpha_{CT} = 4, \alpha_{GT} = 1)$, as three independent vectors (one per codon position)
- Codon equilibrium frequencies $\sim$ Dirichlet$(\alpha_1 = 2, \alpha_2 = 2, \ldots, \alpha_{61} = 2)$

Under these conditions, the mean tree height was 0.46 substitutions per codon position with a 95% HPD of (0.05, 1.05). This translates to 1.38 substitutions per nucleotide site, and a 95% HPD of (0.15, 3.15).

In the P3 + R4 + W4 inference model (nucleotide-based), the following priors were used:

- Speciation rate $\lambda \sim \mathrm{LogNormal}(\mathrm{mean} = 1, \sigma = 2)$
- Reproduction number $\frac{\lambda}{\mu} \sim \mathrm{LogNormal}(\mathrm{mean} = 3, \sigma = 0.5)$
- Relaxed clock standard deviation $\sim \mathrm{LogNormal}(\mathrm{mean} = 0.4, \sigma = 0.5)$
- One bModelTest[13] substitution model average framework per codon position
- Codon relative rates $\sim$ Dirichlet$(\alpha_1 = 1, \alpha_2 = 1, \alpha_3 = 1)$
- Free rates – relative rates $\sim$ Dirichlet$(\alpha_1 = 1, \alpha_2 = 1, \alpha_3 = 1, \alpha_4 = 1)$, one vector per codon position
- Free rates – category weights $\sim$ Dirichlet$(\alpha_1 = 4, \alpha_2 = 4, \alpha_3 = 4, \alpha_4 = 4)$, one vector per codon position

#### 4.5 nN/nS mammal studies

To perform inference on our four chosen mammalian loci, the following prior distributions were used in nucleotide-based inference:

- Speciation rate $\lambda \sim \text{LogNormal}(\text{mean} = 1, \sigma = 2)$
- Reproduction number $\frac{\lambda}{\mu} \sim \text{LogNormal}(\text{mean} = 3, \sigma = 0.5)$
- Gamma spike clock standard deviation $\sim \text{LogNormal}(\text{mean} = 0.4, \sigma = 0.5)$
- One bModelTest[13] substitution model average framework per codon position
- Codon relative rates $\sim \text{Dirichlet}(\alpha_1 = 1, \alpha_2 = 1, \alpha_3 = 1)$
- Free rates – relative rates $\sim \text{Dirichlet}(\alpha_1 = 1, \alpha_2 = 1, \alpha_3 = 1, \alpha_4 = 1)$, one vector per codon position
- Free rates – category weights $\sim \text{Dirichlet}(\alpha_1 = 10, \alpha_2 = 10, \alpha_3 = 10, \alpha_4 = 10)$, one vector per codon position

The following priors were used in codon-based inference (also using free rates for site heterogeneity, and the same tree and clock models):

- Speciation rate $\lambda \sim \text{LogNormal}(\text{mean} = 10, \sigma = 0.5)$
- Reproduction number $\frac{\lambda}{\mu} \sim \text{LogNormal}(\text{mean} = 4, \sigma = 0.5)$
- Gamma spike clock standard deviation $\sim \text{LogNormal}(\text{mean} = 0.4, \sigma = 0.5)$
- $\omega \sim \text{LogNormal}(\text{mean} = 1, \sigma = 1)$
- Transition-transversion ratio $\sim \text{LogNormal}(\text{mean} = 2, \sigma = 1)$
- Codon equilibrium frequencies $\sim \text{Dirichlet}(\alpha_1 = 2, \alpha_2 = 2, \ldots, \alpha_{61} = 2)$
- Free rates – relative rates $\sim \text{Dirichlet}(\alpha_1 = 1, \alpha_2 = 1, \alpha_3 = 1, \alpha_4 = 1)$
- Free rates – category weights $\sim \text{Dirichlet}(\alpha_1 = 5, \alpha_2 = 5, \alpha_3 = 5, \alpha_4 = 5)$

#### 4.6 Case study 1: mammals

Inference was done under the multispecies coalescent model, powered by StarBeast3.[14] Loci were partitioned into codon positions, with each partition having an independent HKY substitution model with frequencies estimated, and a free rates site heterogeneity model with four categories.

The following priors were used:

- Yule skyline tree prior[3] with 10 epochs and birth rate *rate* $\sim$ LogNormal$(\text{mean} = 1, \sigma = 1)$
- Species mean effective population size $\sim$ LogNormal$(\text{mean} = 0.1, \sigma = 1.25)$
- Species tree relaxed clock standard deviation $\sim$ Gamma$(\alpha = 5, \beta = 0.05)$
- Relative clock rate, per partition $\sim$ LogNormal$(\text{mean} = 1, \sigma = 0.6)$
- Transition-transversion ratio, per partition $\sim$ LogNormal$(\mu = 1, \sigma = 1.25)$
- Nucleotide frequencies, per partition $\sim$ Dirichlet$(\alpha_A = 1, \alpha_C = 1, \alpha_G = 1, \alpha_T = 1)$
- Free rates – relative rates, per partition $\sim$ Dirichlet$(\alpha_1 = 1, \alpha_2 = 1, \alpha_3 = 1, \alpha_4 = 1)$
- Free rates – category weights, per partition $\sim$ Dirichlet$(\alpha_1 = 10, \alpha_2 = 10, \alpha_3 = 10, \alpha_4 = 10)$

#### 4.7 Case study 2: anticodon binding domains

The three data types – amino acids, 3Di, and indels – were assigned own relative clock rate. We used the OBAMA model averaging framework for estimating the amino acid site model.[15] The VT substitution model was chosen with 100% probability.[16] The same model averaging framework was employed for the 3Di model, however the three empirical substitution models being compared were the FoldSeek 3Di matrix[17] and the two matrices by Garg and Hochberg[18] (one trained on AlphaFold data and the other on large language models). This selected the Garg and Hochberg AlphaFold model with 100% posterior support. This 3Di model average package is available at https://github.com/jordandouglas/FoldBeast. Lastly, we used a binary Mk substitution model[19] with four gamma rate categories to model indel evolution.

The following priors were used in our analysis of anticodon binding domains.

- Yule skyline tree prior[3] with 10 epochs and birth rate *rate* $\sim$ LogNormal$(\text{mean} = 1, \sigma = 1)$
- Three independent relaxed clock models (aa, 3Di, and indels) each with a branch rate standard deviation $\sim$ Gamma$(\alpha = 5, \beta = 0.05)$
- aa partition relative clock rate fixed to 1
- 3Di partition relative clock rate $\sim$ LogNormal$(\text{mean} = 0.1, \sigma = 0.5)$
- indel partition relative clock rate $\sim$ LogNormal$(\text{mean} = 0.1, \sigma = 1)$

##### 4.8 Case study 3: influenza

The H3N2 dataset was analysed under the BICEPS tree prior[3] with all default prior settings. We used bModelTest to estimate the site and substitution model.[13]

The following priors were used:

- Strict clock molecular rate $\sim$ LogNormal$(\text{mean} = 0.005, \sigma = 0.3)$ substitutions per site per year
- Strict clock deme migration rate $\sim$ LogNormal$(\text{mean} = 0.5, \sigma = 0.5)$ changes per year
- Relative rates among the 4 demes $\sim$ Gamma$(\alpha = 1, \beta = 1)$ (one rate per pairwise transition)
- Deme relative rate indicators: total sum - 3 $\sim$ Poisson$(\mu = 0.693)$, to a maximum sum of 6

### 5 Biological dataset preparation and analysis

#### 5.1 Case study 1: mammals

We downloaded all loci from OrthoMaM v12.[20] These curated alignments describe coding regions. To select our 50 loci for phylogenetic analysis, first we removed all codon sites that contained any gaps or ambiguous characters. Then, we filtered out all loci that were not represented by all 190 species. After these steps, we selected the 50 longest loci, which ranged from 816 – 4983 nucleotides in length (and therefore 272 – 1661 codons).

We performed a phylogenetic independent contrast method described by Welch and Waxman.[21] Taxon pairs were selected automatically using a window approach, such that each sampled pair is a) phylogenetically independent, meaning that their paths do not intersect in the phylogeny, and b) the two taxa share a common ancestor with an age between 0.01 and 0.03 substitutions per site in the time tree. This algorithm was run on 10,000 samples of time tree from the posterior distribution.

In each replicate, linear regression is performed on the differences between the two taxon values $\Delta x$ and $\Delta y$. $\Delta x$ is defined as the difference in adult body mass between two species. $x$ is first logged and standardised such that the term has a mean of 0 and standard deviation of 1. $\Delta y$ is the difference in the natural logarithm of the number of synonymous changes to the most-recent common ancestor of the two taxa. To reduce the influence of extreme values and outliers, any coordinates of $\Delta x$ and $\Delta y$ were removed if they reside more than three standard deviations from the mean of either term. A negative correlation between $\Delta x$ and $\Delta y$ can therefore be interpreted as a negative association between body size and mutation rate.

#### 5.2 Case study 2: anticodon binding domains

To build our alignment of 143 anticodon binding domains, we aligned the atomic coordinate files of this domain from AARS Online.[22] Amino acid and 3Di alignments were generated by FoldMason,[23] followed by a manual removal of poorly aligned structures and regions that contained many gaps.

#### 5.3 Case study 3: influenza

We summarised the posterior distribution of influenza trees using a new BEAST 2 tool called SegmentedTreeAnnotator. First, the trees are logged as segmented trees, such that every state change in the tree (i.e., location changes in this case) is associated with a unary node. This means that branches are broken up into one or more segments, where each segment has a parent that is either a) a binary node describing a bifurcation event, or b) a unary node that has a different sequence/state. To summarise this posterior distribution, we first generate a summary tree from the unsegmented tree (the CCD0 tree in this case) and then split its branches into segments based on the posterior distribution of segmented trees. Given a clade $C$ in the summary tree, the segments along the branch above $C$ describe the *maximum a posteriori* substitution pathway along that branch. The times of those segments is then the mean or median time in the posterior averaged across the clade (as well as 95% HPD). This approach can give negative segment lengths, and therefore we offer a third option – uniform – which staggers the segment times uniformly along the branch to aid with visualisation purposes (which we did in the main article). Segmented trees are saved as nexus files and can be readily displayed using programs like IcyTree.[24]
